# Organoid Modeling and Single-Cell Profiling Uncover the Migration Mechanism of Smooth Muscle Cells in Moyamoya Disease

**DOI:** 10.1101/2024.09.01.610617

**Authors:** Shihao He, Junze Zhang, Xilong Wang, Zhen Qi, Zhenyu Zhou, Yanru Wang, Shaoqi Xu, Dandan Li, Xun Ye, Ziqi Liu, Xiaokuan Hao, Yuanli Zhao, Rong Wang

## Abstract

Moyamoya disease (MMD) is a chronic cerebrovascular disorder characterized by progressive occlusion of the intracranial arteries, resulting in severe ischemic or hemorrhagic stroke. The main characteristic of the affected vessels in MMD is arterial intimal thickening. However, there are no in vitro or in vivo models that can mimic its vascular characteristics. Moreover, the mechanisms underlying the intimal thickening remain unexplained. Here, we generated vascular organoids by differentiating human induced pluripotent stem cell derived from the peripheral blood of MMD patients, thereby creating an organoid model reflecting both the genetic background and characteristics of the affected vessels. Through single-cell sequencing, we found the increased vascular smooth muscle cell (VSMC) proportion and its functional abnormalities in MMD organoids. Proteomics and RNA sequencing identified abnormal TUBA4A and TUBB4B overexpression in both the organoids and patient serum. The following in vitro experiments demonstrated that TUBA4A and TUBB4B promote the contractile-to-synthetic phenotypic switching, migration and proliferation in VSMC. Further experiments identified the GJA1-mediated PI3K/AKT/KLF4 pathway as a key regulating pathway of these phenotypic changes in VSMCs. Our findings demonstrate that the abnormal expression of TUBA4A and TUBB4B in VSMC might be a significant contributor to the intimal thickening in the affected vessels of MMD.

## Introduction

Moyamoya disease (MMD) is a chronic cerebrovascular disorder characterized by the neuroimaging findings of progressive narrowing or occlusion of the terminal segments of the internal carotid arteries (ICAs) and abnormal neovascularization at the skull base which causes cerebral hemodynamic disturbances, resulting in cognitive impairment, muscle weakness, and even stroke^1^.

Although pathologic studies of MMD have shown that thickening caused by vascular smooth muscle cell proliferation and migration have been detected in the intima of terminal segments of internal carotid arteries, leading to narrowing of the vessel lumen and *RNF213* has been identified as the first susceptibility gene, research on the pathogenesis underlying MMD has remained virtually stagnant for a long time^2–5^.

Appropriate research models can greatly assist in the study of the unknown etiology. Unfortunately, previous efforts to construct research models of MMD have ended in failure^6^. The surgical models dedicated to simulating chronic intracranial ischemia due to MMD by surgical ligation or narrowing of the common carotid artery (CCA) or ICA in animals were limited by poor reproducibility, susceptibility to unexpected acute ischemic lesions, and absent of gene factors, which resulted in a failure to recapitulate the characteristic smooth muscle cell proliferation of MMD^6–8^. While the genetic models in which the *RNF213* was knock-out or knock-in in mice also failed to lead to anticipated histopathological changes^6,9,10^.

Organoids, the in vitro 3D tissues obtained by stem cell culture and differentiation, have the capability to exhibit the anatomical and functional characteristics of real organs, and have been applied to construct disease models, organ development models, and other fields^11^. Commonly, organoids derived from human adult stem cells (ASCs) or induced pluripotent stem cells (iPSCs) can present features relevant to human diseases, and thus be applied as models in the study of disease pathogenesis^11^. So far, a variety of organoid-based disease models have been developed for genetic diseases, host-pathogen interactions, or cancers, which appropriately demonstrate known pathological characteristics^11–14^.

To investigate the etiology of MMD, we have built a model based on vascular organoids derived from iPSCs of MMD patients, identified the response ability of this organoid-based model to the hallmarks of MMD, and further found that overexpression of tubulin alpha (TUBA) might be a novel target for the proliferation and migration of vascular smooth muscle cells in MMD.

## Methods

### Study participants and samples

Patients examined between January 2020 and September 2023 with moyamoya disease (MMD) were clinically diagnosed by medical history and neuroradiological examination following the diagnostic guidelines^15^. Patients diagnosed with MMD (MMD) were classified into two subgroups: adult patients with hemorrhagic history (HEM) and adult patients with ischemic history (IS). Healthy adult volunteers without MMD or any underlying diseases were enrolled as controls (HC). The clinical data of all the patients and controls were recorded. Clinical and imaging data were obtained from medical records and Picture Archiving and Communication Systems, respectively. Written informed consent was obtained from all participants (parents/guardians of patients aged < 18 years). This study was approved by the Institutional Ethics Committee of Beijing Tiantan Hospital, Beijing, China (KY2023-204-02).

Peripheral blood samples for vascular organoids construction were collected from MMD patients and health control (HC) adults by venipuncture using a CPT Vacutainer containing sodium citrate as an anticoagulant prior to revascularization surgery. The subsequent procedures for inducing iPSCs from peripheral blood and constructing organoids are detailed in Section 2.2 of the Methods. Peripheral blood samples for DIA proteomic sequencing were collected from MMD patients and health control (HC) adults as described above. The Vacutainers were centrifuged for 8 min at 460 g. The serum with lymphocytes was transferred into a new Falcon tube and centrifuged again at 3000 r/min for 15 min. The serum supernatant was removed and stored in cryovials at –80℃ prior to use. STA (1-3 mm) samples for the immunofluorescence (IF) staining were obtained during direct or combined revascularization surgery. Blood vessel specimens were shock-frozen in liquid nitrogen prior to RNA extraction. STA tissues were immediately immersed in 4% formalin for subsequent IF staining.

### Vascular organoids generation

Peripheral venous blood (8 mL) from Moyamoya disease patients was collected into anticoagulant tubes and maintained at 37°C. PBMCs were isolated using Ficoll density gradient centrifugation. Whole blood was diluted 1:1 with saline, gently mixed, and layered onto the separation medium (separation medium: diluted blood = 1:2). After centrifugation at 800g for 20-30 minutes at room temperature, the PBMC layer was carefully aspirated and transferred to a 15 mL centrifuge tube. Cells were resuspended in 10 mL of dilution buffer and centrifuged at 250g for 10 minutes at room temperature. This washing step was repeated 1-2 times.

PBMC derived from peripheral blood were reprogrammed into iPSC lines using the CytoTune 2.0 iPS Sendai Virus Reprogramming Kit (ThermoFisher). On Day −4, peripheral blood mononuclear cells (PBMCs) were seeded at a density of 5×10⁵ cells/mL in the center of a 24-well plate with complete PBMC culture medium. From Day −3 to Day −1, half of the medium was replaced daily with 0.5 mL of fresh PBMC culture medium. On Day 0, cells were transduced with CytoTune™ 2.0 Sendai reprogramming vectors and incubated overnight. The following day, the medium was replaced to remove the vectors. On Day 3, the transduced cells were plated on rhVTN-N coated dishes in StemPro™-34 medium without cytokines. From Day 4 to Day 6, the medium was replaced every other day. On Day 7, the transition to iPSC medium began by replacing half of the StemPro™-34 medium with complete iPSC medium. On Day 8, the medium was fully replaced with iPSC medium, and the cells were cultured on Matrigel-coated dishes. From Day 9 to Day 28, the medium was replaced daily, and iPSC colonies were monitored. Once iPSC colonies were ready, live staining was performed, and undifferentiated iPSCs were picked and transferred to fresh Matrigel-coated dishes for expansion and purification.

IPSCs were differentiated into vascular organoids following the methods reported in previous study^16^. IPSCs were first cultured in mTeSR1 medium on Matrigel-coated plates until they reached 80-90% confluency. The cells were then dissociated into single cells using Accutase, resuspended in aggregation medium, and seeded at a density of 2×10^5 cells per well in low-attachment 6-well plates containing 50 μM Y-27632. The cells were incubated for 1 day to form smooth aggregates. On Day 0, the aggregates were transferred to a new medium (N2B27 Medium supplemented with 12 μM CHIR99021 and 30 ng/mL BMP-4) and cultured for 3 days to induce mesoderm differentiation. The medium was gently pipetted daily to prevent clumping. On Day 3, the aggregates were transferred to N2B27 Medium supplemented with 100 ng/mL VEGF-A and 2 µM forskolin and cultured for 2 days. The cell aggregates were then embedded in a two-layer Collagen I-Matrigel mixture in 12-well plate (0.5 mL for layer 1 and 0.5 mL for layer 2) and incubated at 37°C to solidify each layer. After embedding, StemPro-34 SFM medium with 15% FBS, 100 ng/mL VEGF-A, and 100 ng/mL FGF-2 was added. The organoids were cultured for 1-3 days, with medium changes every other day, to induce vascular differentiation and sprouting. At day 10, the vascular networks were formed and extracted from the Collagen I-Matrigel to a 96-well ultra-low-attachment plate to self-assembly form VOs.

### Transcriptome Sequencing

For each sample, 10 organoids were harvested and washed with PBS to remove residual culture media. RNA was extracted from organoid samples using an RNAprep Pure Micro Kit (DP420, Tiangen Biotech). The libraries were generated according to the manufacturer’s protocol of NEBNext Ultra Directional RNA Library Prep Kit for Illumina. Then the sequencing was carried out on Illumina NovaSeq 6000 platform. After sequencing error rate distribution checks, GC content distribution checks, and raw data filtering, clean reads were obtained for each sample. The identified DEGs required an FDR value < 0.01 and a log2FC > 1. The RNA-seq analysis was performed by Novogene Bioinformatics Technology Co. Ltd. (Beijing, China).

### Single-cell RNA sequencing

For each sample, 20 organoids were collected and washed with cold PBS to remove any residual culture media. The organoids were then dissociated into single cells using enzymatic digestion. Cell viability was assessed using Trypan Blue exclusion, and only samples with a viability exceeding 85% were processed further. For each sample, 10,000 viable single cells were collected for sequencing. The single-cell suspension was then loaded onto a Chromium Single Cell Controller (10x Genomics) to create single-cell gel bead-in-emulsions (GEMs). Subsequent cDNA library construction was carried out using the Single Cell 3’ Library & Gel Bead Kit v3 (10x Genomics) as per the manufacturer’s guidelines. Sequencing was performed on a 10x Genomics platform at Novogene Co., Ltd. (Beijing, China). The sequencing data were then processed and analyzed using standard bioinformatics pipelines to explore gene expression at the single-cell level.

### DIA proteomic sequencing

DIA quantification proteomics was used to identify differences in protein expression between MMD patients and HC. The DIA proteomics detecting methods were mentioned in our previous study^17^. Bioinformatics analysis was performed using the R software (version 3.4, R Foundation for Statistical Computing). The DIA analysis was performed by Beijing Genomics Institute (BGI, Shenzhen, China).

### Enzyme-Linked immunosorbent assay

Enzyme-linked immunosorbent assays (ELISA) were used to measure human TUBA4A and TUBB4B protein in serum samples using Human Tubulin Alpha-4A ELISA Kit (EKL61314, Biomatik, Canada) and Human Tubulin Beta-4B Chain ELISA Kit (Human) (abx548829, Abbexa, USA).

### Cell culture and treatment

Human cerebrovascular smooth muscle cells (HBVSMCs) were cultured. The culture conditions were SMCM medium containing 2% FBS, 1% P/S double antibody and 1% smooth muscle cell growth supplement (SMCGS). The HBVSMC cells were incubated in DMEM containing 2% FBS for 24h, followed by incubation in DMEM medium containing 2.5% heat-inactivated human serum from MMD patients and HC for an additional 24h.

### Plasmid construction and small hairpin RNA transfection

Plasmids encoding TUBA4A, TUBB4B and GJA1 were purchased from the Boen Company (Guangzhou, China). ShRNAs against TUBA4A, TUBB4B and GJA1, and the corresponding controls were obtained from Boen Company (Guangzhou, China). HBVSMCs cells were transfected with various types of lentiviruses at a multiplicity of infection of 10 and 5 µg/ml puromycin was used for selection for 2 weeks to obtain stably transfected cell lines. PCR and western blotting assays were used to verify the transfection efficiency of the lentiviruses. The detailed procedures for plasmid and shRNA construction can be found in the supplementary materials.

### Cell proliferation assay

Cell proliferation abilities of HBVSMCs were evaluated using a Click-iT™ EdU imaging kit (C0085L, Beyotime biotechnology) according the protocols of the manufacturer. The cells were observed and imaged under a fluorescence microscope using red (Ex/Em = 495/519 nm) filters.

### Wound-Healing Assay

Wound-Healing Assay was used to analyze the migration ability of HBVSMCs. The treated HBVSMC cells were seeded into 6-well plates (5 × 10^5^ cells per well) in DMEM containing 10% FBS and maintained at 37°C in 5% CO_2_ overnight. Subsequently, a 200 μl pipette tip was used to create scratches across the cell monolayer, followed by washing the cells three times with PBS. Each well was then supplemented with 2ml of DMEM and incubated for 48h. Microscopic images were captured at 200× magnification.

### Western Blot

The treated cells were resuspended and seeded in 6-well culture plates (5×10^5^ cells per well). After the cells were attached to the wall, whole-cell lysates were prepared using RIPA buffer. The protein concentration was determined using a Pierce BCA Protein Assay kit (Thermo Fisher Scientific). After electrophoresis, electrotransfer, and incubation with the primary and secondary antibodies, signals were detected using Novex ECL HRP chemiluminescent substrate reagent kit (WP20005, Thermo Fisher Scientific). Glyceraldehyde 3-phosphate dehydrogenase (GAPDH) was used as the loading control. The primary antibodies and their concentration used were listed in the supplementary materials (Table S2).

### Flow cytometric cell cycle analysis

The treated HBVSMC cells were seeded into 6-well plates (5 × 10^5^ cells per well) in DMEM containing 10% FBS and maintained at 37°C in 5% CO_2_ overnight. The cells were collected, washed with PBS, and the concentration was adjusted to 1×10^6^ cells/ml. Single-cell suspensions of cells were fixed in 70% ethanol at 4℃ overnight.

The cell cycle stage of HBVSMC cells was determined using a cell cycle assay kit (KGA512, KeyGEN Biotech, NanJing, China). The samples were then analyzed using a flow cytometer (Beckman DxFlex, Beckman) at 488 nm excitation. The data were analyzed using FlowJo software (FlowJo, LLC, Ashland, OR, USA). The percentages of cells in the G1, S, and G2 phases were calculated, respectively.

### Immunofluorescence staining

Immunofluorescence (IF) staining was performed on vascular organoids following the protocol previously described in the literature^16^. As for STA tissue samples, STA tissue samples were fixed in 4% paraformaldehyde at room temperature for 24 hours, followed by embedding in paraffin and sectioning into 4 µm thick slices. Sections were deparaffinized in xylene and rehydrated through graded alcohols. Antigen retrieval was performed by heating the sections in citrate buffer (pH 6.0) at 95°C for 20 minutes. After cooling to room temperature, sections were blocked with 5% bovine serum albumin (BSA) in PBS for 1 hour to prevent non-specific binding. The tissue sections were then incubated with fluorophore-conjugated primary antibodies at 4°C overnight. The following day, sections were washed three times with PBS. After washing, nuclei were counterstained with DAPI, and the sections were mounted with antifade mounting medium. Fluorescent images were captured using a confocal microscope at appropriate magnifications. The antibodies and their concentration used were listed in the supplementary materials (Table S2).

### Statistical analysis

Data were analyzed and visualized using GraphPad Prism 9 (Version 9.4.0) and organized with Adobe Illustrator 2022 (Version 2022). Results are presented as means ± SD, and statistical differences between groups were assessed using one-way ANOVA followed by Tukey’s test. A P value of less than 0.05 was considered statistically significant.

## Results

### Patients and samples

This study included 105 patients diagnosed with MMD between January 2020 and August 2022. Peripheral blood samples were taken from all 76 patients. Six patients and 3 health control (HC) individuals were selected from whom peripheral blood was used to generate iPSCs and subsequently construct vascular organoids. DIA quantification proteomics were performed on the peripheral blood of 20 health controls (HC) individuals and 40 MMD patients (20 HEM and 20 IS), including 20 ischemic (IS) and 20 hemorrhagic (HEM) MMD. In the validation group, we first used ELISA to validate the candidate proteins found in the DIA data of 20 peripheral blood samples (random selection of 5 from each group in DIA samples; IS = 5, HEM = 5 and HC = 5). We then used ELISA experiment to validate the proteins in the peripheral blood of 60 samples (another 15 were randomly selected for each group; IS = 15, HEM = 15 and HC = 15). Serum extracted from 13 peripheral blood samples from DIA experiments was used for the in vitro cell experiments. Detailed information on all patients can be found in the supplementary materials. (Table S1).

### Generation of vascular organoid model for HC and MMD

We collected the peripheral blood from 3 MMD patients and 3 HC. According to the previous protocols^18,19^, the peripheral blood monocular cells (PBMCs) were extracted from peripheral blood and then were reprogrammed into induced pluripotent stem cells (iPSCs). We subsequently differentiated the reprogrammed iPSCs from MMD patients and HC into vascular organoids, following the methods reported in previous study^16^. The overview of the process for developing the vascular organoids was shown in **Figure 1A**. **Figure 1B** shows the morphological progression of the various stages from iPSC aggregates to vascular organoids under light microscopy.

**Figure 1:**
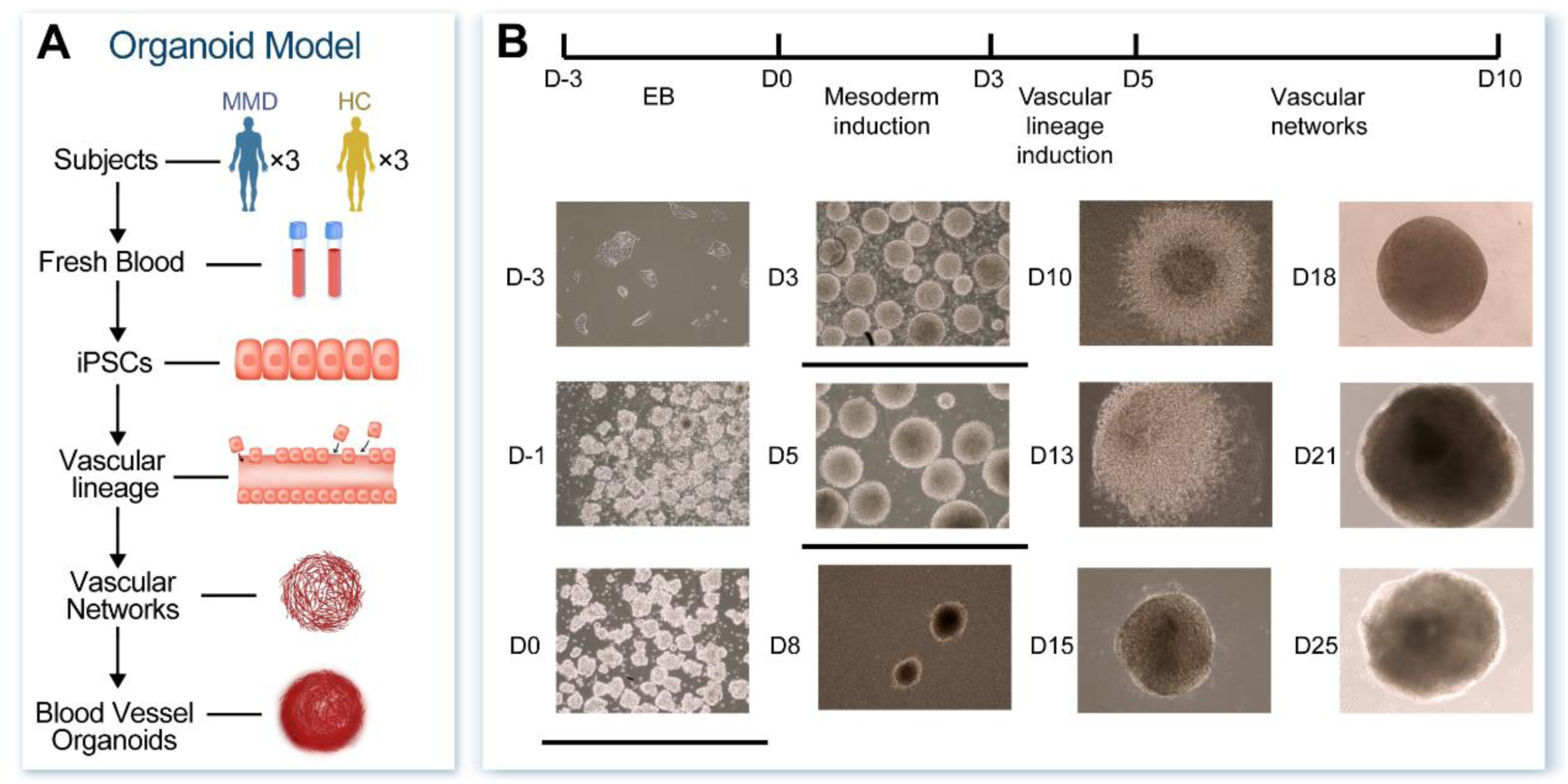
Generation of the vascular organoids derived from HC individuals and MMD patients. **A,** The flowchart of vascular organoid generation derived from HC and MMD patients. **B,** Bright-field images of human induced pluripotent stem cells (iPSCs) differentiation into vascular organoids (day-3 – day25). Scale = 300μm. MMD = Moyamoya Disease; HC = Health Control.

To verify the morphology of the vascular network and organoids, we performed immunofluorescence (IF) staining on the differentiated vascular network at day 10 and vascular organoid at day 25. Vascular networks at day 10 exhibit a highly branched and connected endothelial (CD31+) and smooth muscle cell (αSMA+) network (**Figure 2A, 4A**). The 3D reconstruction of the vascular organoids demonstrates the spatial morphology of the vascular organoids, and an interweaving of endothelial cells (CD31+) and smooth muscle cells (αSMA+) within the organoids (**Figure 3A**). The vascular network in vascular organoids derived from MMD patients was more extensive than that form HCs. It suggests an excessive vascular proliferation in the vascular organoids derived from MMD patients.

**Figure 2:**
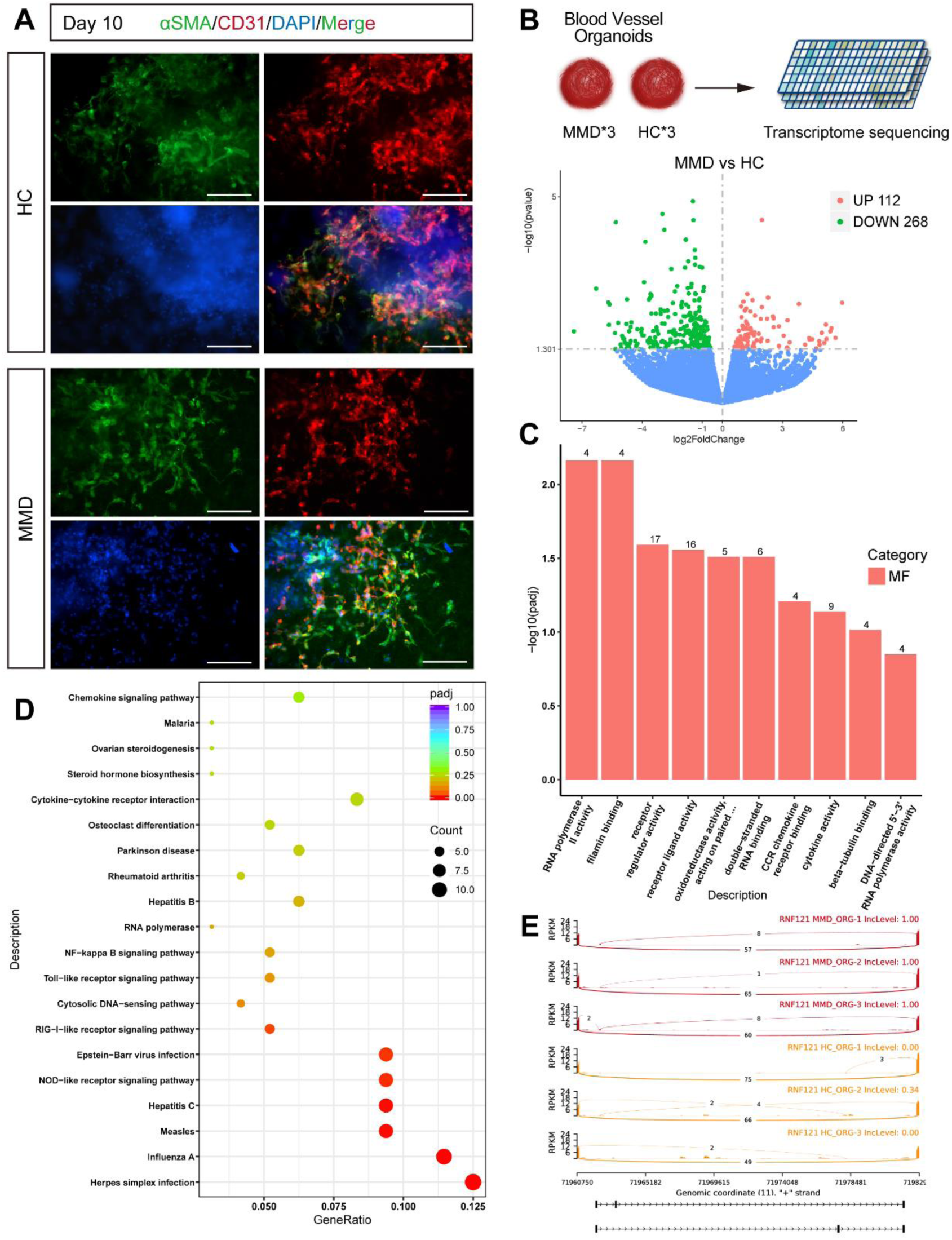
Transcriptome sequencing of vascular organoids suggests that the abnormal gene expression in the vascular organoids. **A,** Representative immunofluorescence of CD31 and αSMA expressing endothelial and smooth muscle cells respectively shows establishment of vascular networks. CD31, red; αSMA, green; DAPI, blue; HC, health control; MMD, moyamoya disease. Scale = 100μm. **B,** Volcano plot showing the DEGs between vascular organoids derived from MMD patients and HC. **C,** MF enrichment analysis of DEGs between vascular organoids derived from MMD patients and HC. **D,** KEGG enrichment analysis of DEGs between vascular organoids derived from MMD patients and HC. **E,** The alternative splicing analysis of vascular organoid. KEGG = Kyoto Encyclopedia of Genes and Genomes; DEGs = differentially expressed genes; MF = Molecular function; ORG = Organoid.

**Figure 3:**
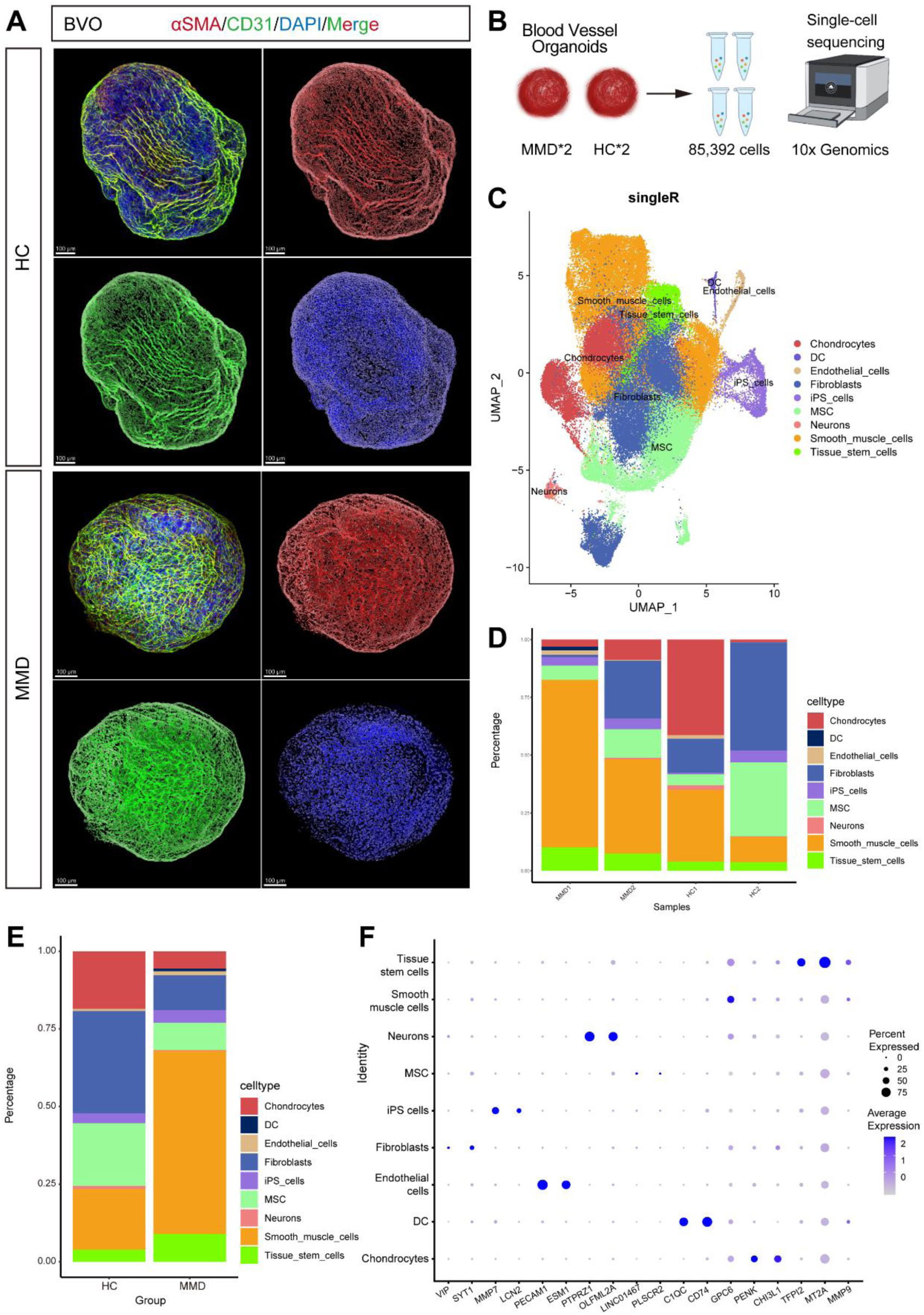
Single-cell RNA sequencing and histological staining indicate an elevated number of smooth muscle cells in MMD vascular organoids and arterial wall. **A,** Representative immunofluorescence and three-dimensional (3D) reconstruction of HC and MMD vascular organoids. CD31, green; αSMA, red; DAPI, blue. Scale = 100μm. **B,** Schematic representation of sc-RNA sequencing. A total of 85,392 single cells were captured, including organoids derived from 2 MMD patients and 2 HC. **C,** A UMAP plot displaying all cells colored according to the 9 major cell types. **D,** The proportion of different cell clusters within each organoid. **E,** The percentage of each cell type in vascular organoids derived from MMD and HC. **F,** The main gene expression profiles of different cell types in MMD and HC vascular organoids. Sc-RNA = Single cell RNA sequencing.

### Transcriptome sequencing of vascular organoids suggests that the abnormal gene expression in the vascular organoids

We performed bulk RNA sequencing on organoid tissues to identify differentially expressed genes (DEGs) between organoids derived from MMD patients and HC individuals (**Figure 2B**). Compared to organoids from HC, vascular organoids derived from MMD patients had 112 genes upregulated and 268 genes downregulated (**Figure 2B**). GO enrichment analysis for these DEGs revealed significant enrichment in molecular functions (MF) related to beta tubulin binding, filamin protein binding, etc. (**Figure 2C**). Kyoto Encyclopedia of Genes and Genomes (KEGG) pathway analysis identified enrichment in pathways such as Toll-like receptor signaling pathway, Cytokine-cytokine receptor interaction, etc. (**Figure 2D**). The alternative splicing analysis of six vascular organoid is shown in **Figure 2E**.

To explore the different proteomics expression patterns between MMD and HC, we systematically analyzed the DIA proteomics data of the serum from 40 MMD patients and 20 HC reported in our previous study^17^. Differential protein analysis revealed a significant upregulation of TUBA4A (Log_2_FC (MMD/HC) =2.7, p < 0.05) and TUBB4B (Log_2_FC (MMD/HC) =2.1, p < 0.05) proteins in MMD patients’ serum (**Figure S2A, S2B**). The chordal graph shows that the TUBA4A and TUBB4B have strong correlation with the structural constituent of cytoskeleton and cadherin binding, which are involved in cell cytoskeleton (**Figure S2C**). Subsequently, the ELISA was performed to validated the upregulation of these two proteins in an additional cohort of 45 individuals’ serum (30 MMD vs. 15 HC) (**Figure 5A-5C**).

The IF staining was performed on the vascular organoids at day 25 and observed an enhanced fluorescence signal of TUBA4A protein in the MMD organoids comparing with HC (**Figure 4A**). The western blot was performed to validate the expression of TUBA4A and TUBB4B in vascular organoids, and found that the two proteins were upregulated in MMD vascular organoids (**Figure 4D**). The vascular organoids and DIA proteomic sequencing both showed the upregulation of TUBA4A and TUBB4B, which may have a correlation with MMD pathogenesis. Additionally, in the vascular organoids, we observed a decrease in the expression levels of smooth muscle contractile markers (CNN1 and SM22α) and an increase in synthetic markers (S100A4 and ZO-2) (**Figure S1A-S1E**).

**Figure 4:**
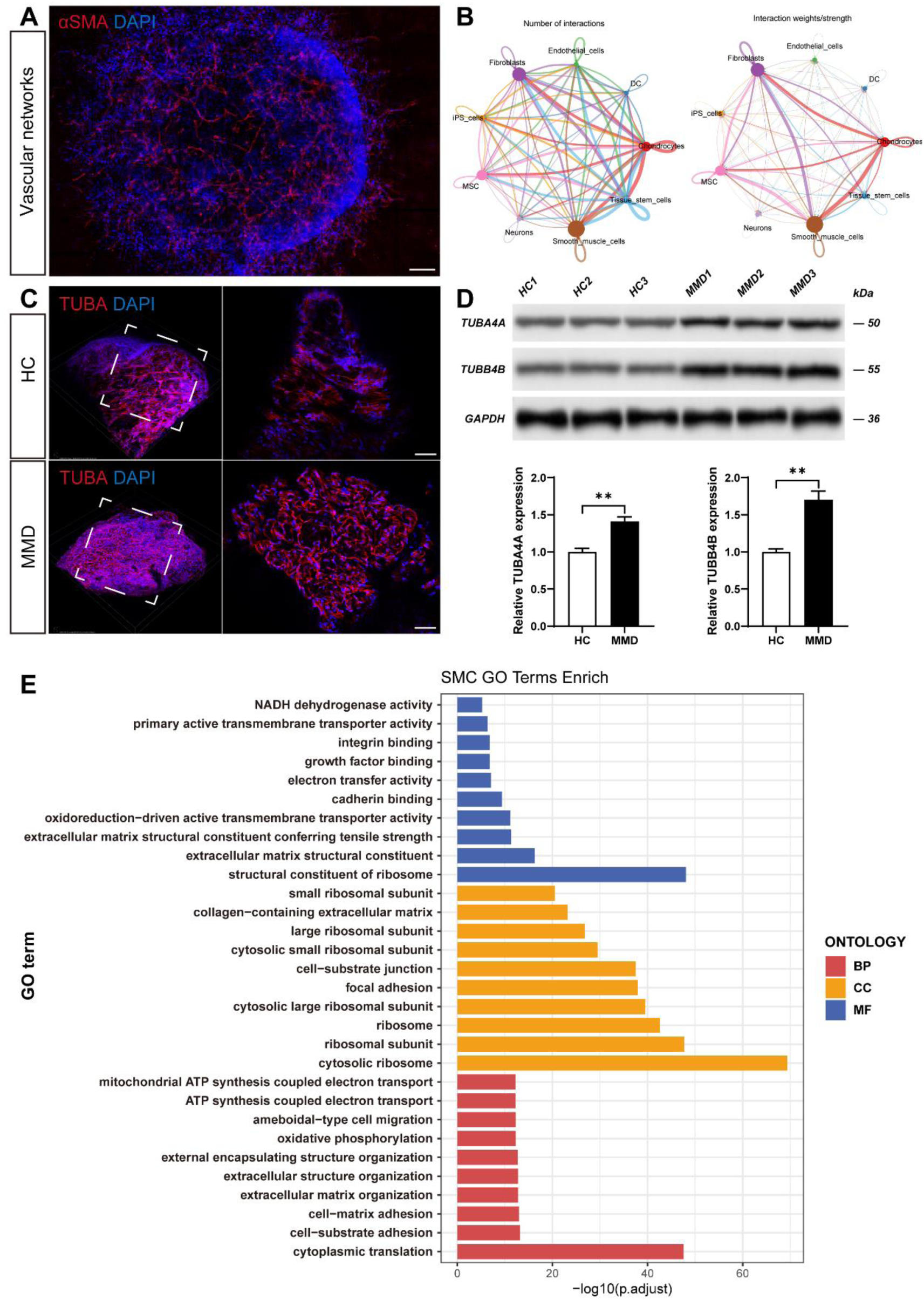
Characteristics of smooth muscle cells in MMD vascular organoids. **A,** Immunofluorescence staining of smooth muscle cells during the vascular network phase in MMD organoids. **B,** The weight and number of connections between smooth muscle cells and other cell types in the MMD organoids. **C,** Immunofluorescence staining of TUBA4A in vascular organoids from MMD patients and HC. TUBA, red; DAPI, blue. **D,** Western blot showing the expression level of TUBA4A and TUBB4B in the vascular organoids. The bar chart showing the expression level of TUBA4A and TUBB4B. **E,** GO enrichment analysis of smooth muscle cells in vascular organoids. GO = Gene Ontology.

### Single-cell RNA sequencing and histological staining indicate an elevated number of smooth muscle cells in MMD vascular organoids and arterial wall

To analyze the cell types of vascular organoids, we performed single-cell RNA sequencing (scRNA-seq) for 2 MMD organoids and 2 HC organoids. By digesting the vascular organoids in day 25, we collected a total of 85,392 single cells followed by scRNA-seq (**Figure 3B**). A total of 9 cell types are annotated, including smooth muscle cells (SMC), endothelial cells, iPSCs, MSC, fibroblasts, DC, neurons, tissue stem cells and chondrocytes (**Figure 3C**). The proportion of various cell type in each sample was shown in **Figure 3D**. By comparing the proportions of different cell types in MMD and HC vascular organoids, we found that the proportion of SMCs in MMD organoids was significantly higher than that in HC organoids (**Figure 3E**). **Figure 3F** shows the main gene expression profiles of different cell types in MMD and HC vascular organoids. We further analyzed the interactions between different cell clusters. SMCs exhibited a high number of interactions and stronger interaction weights with other cell types, including endothelial cells, chondrocytes, MSC and fibroblasts (**Figure 4B**). The GO enrichment analysis of smooth muscle cells in organoids is shown in **Figure 4E**.

We then performed immunofluorescence (IF) staining on the superficial temporal artery (STA) from MMD patients and non-MMD controls (**Figure 7A**). The walls of the STA in MMD patients exhibits a significant intimal thickening, in which an abundant presence of VSMCs was observed (**Figure 7B**). The IF staining for STA and the sc-RNA analysis for vascular organoids both demonstrates the over proliferation of VSMCs in the vessel wall of MMD. These results demonstrates that the a VSMCs might be strongly associated with the pathogenesis of MMD.

### The vascular smooth cells exhibit the abilities of migration, proliferation, and a tendency of contractile-to-synthetic phenotypic switching in MMD

Serum collected from patients in the MMD and HC groups was utilized to incubate HBVSMCs to explore the effect of MMD patient (IS and HEM patients) serum on VSMC (**Figure S3A**). After 24 hours of serum incubation, the MMD serum-treated HBVSMCs showed significant promoted abilities of migration, proliferation and contractile-to-synthetic phenotypic switching compared with that of HC serum-treated HBVSMCs (**Figure 5E, S3B-D, S3G-H**). Interestingly, we also find the upregulation of TUBA4A and TUBB4B protein in the MMD serum-treated HBVSMC (**Figure S3E**). Detailed descriptions of the serum co-culture results can be found in the supplementary materials (**Figure S3**).

**Figure 5:**
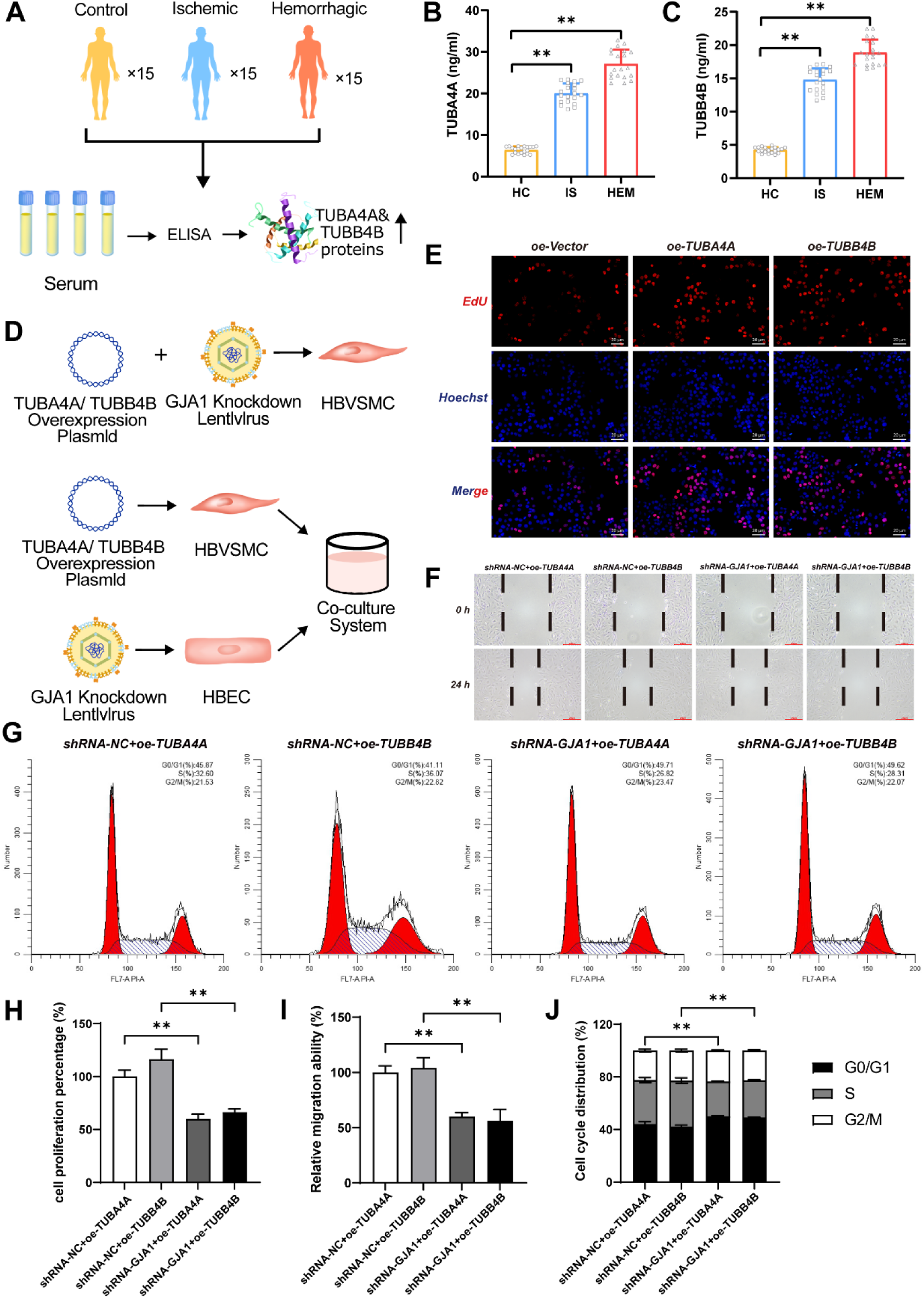
TUBA4A and TUBB4B promote the abilities of migration, proliferation and contractile-to-synthetic phenotypic switching of SMC in MMD. **A,** ELISA validation of differentially expressed proteins identified in serum proteomics. **B,** The bar chart showing the expression levels of TUBA4A in the serum validation cohort. **C,** The bar chart showing the expression levels of TUBB4B in the serum validation cohort. **D,** Schematic diagram of the cell model construction for TUBA4A/TUBB4B overexpression and GJA1 knockdown in HBVSMCs, and the co-culture of smooth muscle cell (SMC) and endothelial cell (EC). **E,** 5-Ethynyl-2′-deoxyuridine (EDU) staining of the TUBA4A/TUBB4B overexpression HBVSMC. Colors: EDU (red), 4′,6-diamidino-2-phenylindole (DAPI) (blue). Scale = 20 µm. The bar chart of **H** shows the proportion of proliferation HBVSMC. **F,** Scratch assay showing the migration ability of GJA1 knockdown HBVSMC. Scale = 200µm. The bar chart of **I** shows the proportion of migration HBVSMC. **G,** Flow cytometric cell cycle analysis for the TUBA4B/TUBB4B overexpression HBVSMC after GJA1 knockdown. The bar chart of **J** shows the proportion of HBVSMC in G0/G1, S and G2/M cell cycle phases. HBVSMC = Human Brain Vascular Smooth Muscle Cells.

To further investigated the role of TUBB4A and TUBB4B in these changes of HBVSMCs, we performed overexpression and knockdown for these two genes using overexpressing plasmid and small hairpin RNAs (shRNA) in HBVSMCs, respectively (**Figure S5A**). The HBVSMCs present high abilities of migration and proliferation, and a tendency of contractile-to-synthetic phenotypic switching in the TUBA4A and TUBB4B overexpressed group (**Figure S4B-C, S5B-D, S5F-G**). Conversely, when knockdown the two genes, the changes of HBVSMCs were reversed in the MMD serum-treated groups (**Figure S4B-C, S5B-D, S5F-G**). Detailed descriptions of the serum co-culture results can be found in the supplementary materials (**Figure S4, S5**). These results suggest that TUBA4A and TUBB4B are essential for these changes of HBVSMCs in MMD, including proliferation, migration, and phenotypic switching.

### TUBA4A and TUBB4B induce the changes of VSMC in MMD via GJA1/PI3K/AKT/KLF4 pathway

When treating HBVSMCs with MMD serum and overexpressing the TUBA4A and TUBB4B, we observed an increased expression of GJA1 and the following PI3K/AKT/KLF4 pathway proteins in HBVSMCs as well as in MMD-derived vascular organoids (**Figure S3F, S5E, S1A, S1F-K**). To study whether GJA1 contributes to these changes in HBVSMCs, we utilized overexpression plasmids and shRNA to upregulate and downregulate the level of GJA1, respectively (**Figure S6A**). After the GJA1 overexpression, the HBVSMCs show enhanced the migration ability, increased proportion of proliferating cells and tendency of contractile-to-synthetic phenotypic switching (**Figure S6C-F**). Conversely, when GJA1 was knocked down in HBVSMCs overexpressing TUBA4A and TUBB4B, we observed these TUBA4A or TUBB4B-induced HBVSMCs changes were reversed (**Figure 5D, 5F, 5G, 5I, 5J**).

To further investigate whether the PI3K/AKT/KLF4 pathway contributes to the GJA1 overexpressed-caused changes of HBVSMCs, we used Kenpaullone, a KLF4 inhibitor, to downregulate the expression of the downstream KLF4 molecule in this pathway (**Figure S6A**). After the KLF4 inhibition for HBVSMCs, the enhanced proliferation, migration, and contractile-to-synthetic phenotypic switching induced by GJA1 overexpression were all blocked (**Figure S6B-F**). These demonstrate that the GJA1/PI3K/AKT/KLF4 pathway contributes to the TUBA4A and TUBB4B induced changes of HBVSMC (**Figure 7C**).

### 3.7 TUBA4A or TUBB4B overexpressed HBVSMCs promote the proliferation of HBECs through KLF4 molecule

Meanwhile, we utilized the co-cultured model of HBVSMC and Human brain endothelial cells (HBEC) to study the to investigate the effects of the TUBA4A/TUBB4B overexpressed VSMC on ECs. (**Figure 5D**). Overexpression of TUBA4A in HBVSMCs promotes the proliferation of the co-cultured HBECs (**Figure 6A, 6B, 6C**). When GJA1 was knocked down in HBECs, the enhanced proliferative ability of HBECs induced by TUBA4A/TUBB4B overexpressed HBVSMCs was decreased (**Figure 6A, 6B, 6C, 6D, 6F**). The proliferative ability of HBECs also decreased after decreasing the expressing level of KLF4 by using a KLF4 inhibitor Kenpaullone to the SMC-EC co-culture system (**Figure 6A, 6B, 6G**). In addition, we also found that the expression level of KLF4 increased in HBECs after co-cultured with TUBA-overexpressing HBVSCMs (**Figure E, G**). However, when GJA1 was knocked down in HBECs, we observed a significant decrease in the KLF4 levels in the co-cultured HBECs (**Figure E, G**). These SMC-EC co-culture results indicate that the elevated levels of KLF4 in TUBA4A/TUBB4B overexpressing HBVSMCs can transfer to HBECs through intercellular GJA1, leading to the excessive proliferation of HBECs (**Figure 7C**).

**Figure 6:**
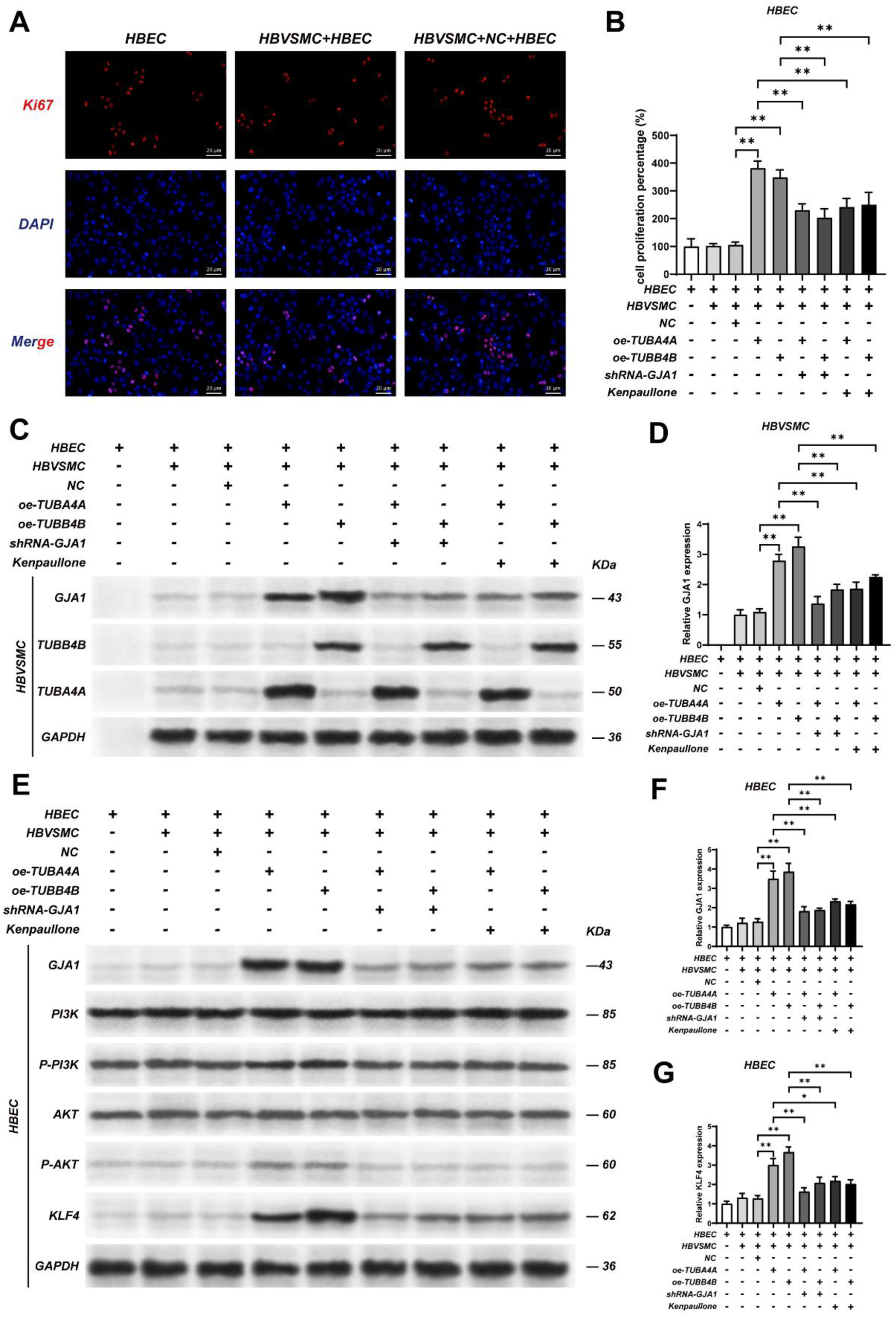
TUBA4A or TUBB4B overexpressed HBVSMCs promote the proliferation of HBECs through KLF4 molecule. **A,** Ki67 staining of Human Brain Endothelial Cells (HBECs) after co-culture with treated HBVSMCs. The bar chart of **B** shows the proportion of proliferation HBEC. **C,** Western blot showing the expression levels of TUBA4A, TUBB4B, and GJA1 in HBVSMCs under different treatment conditions in the SMC-EC co-culture system. The bar chart of **D** shows the expression level of GJA1 in HBVSMCs. **E,** Western blot showing the expression levels of TUBA4A, TUBB4B, and GJA1 in HBECs under different treatment conditions in the SMC-EC co-culture system. The bar chart of **F** and **G** shows the expression level of GJA1 and KLF4 in HBECs, respectively.

**Figure 7:**
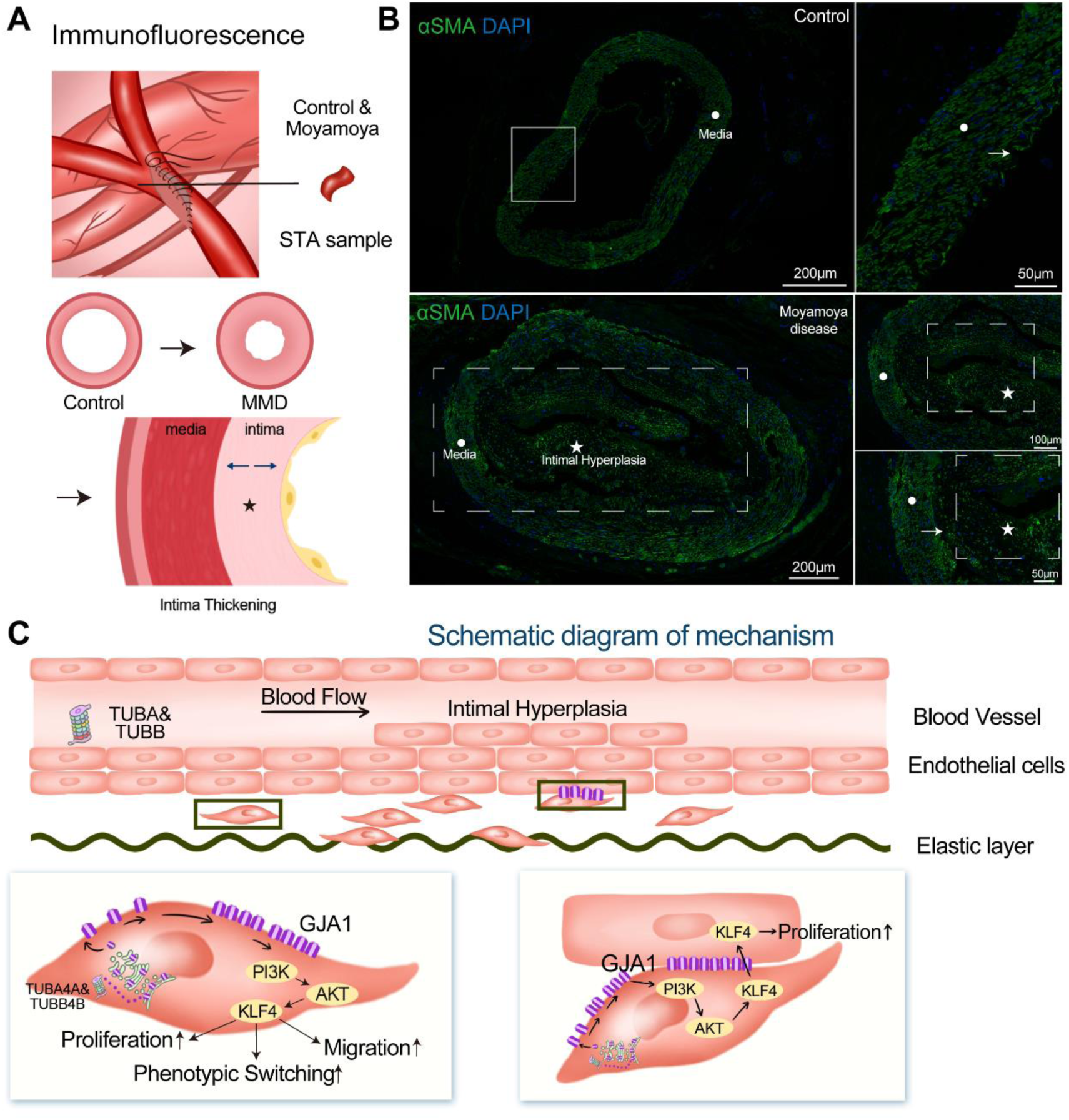
Immunofluorescence staining of intimal thickening in MMD vessels and the underlying mechanism. **A.** Schematic diagram illustrating the collection of STA specimens during surgery and the intimal thickening in MMD arteries. **B,** Immunofluorescence staining of the STA in control subjects and MMD patients. Scale = 200μm. αSMA, green; DAPI, blue. **C,** Schematic diagram of the underlying mechanism for intimal thickening in MMD. STA = Superficial Temporal Artery.

## Discussion

Moyamoya disease (MMD) can seriously affect patients’ quality of life and even result in severe ischemic and hemorrhage strokes^20^. However, there is a lack of effective diagnostic and treatment strategies for MMD because of the unknown pathogenesis underlying it^20^.There is an urgent need for suitable methodologies to further elucidate the mechanisms underlying MMD. Previously, endeavors undertaken by scholars to establish research models pertaining to MMD have ended in failure^6^. Here we reported the construction of the first organoid model of MMD which may significantly advance the progress of research on the pathophysiology of MMD. Herein, we presented the development of the initial organoid model for MMD, which might be anticipated to markedly enhance the advancement of scientific inquiry into the disease’s pathophysiological mechanisms.

Chronic ischemia, a consequence of progressive stenosis in the intracranial segments of the bilateral internal carotid arteries, is a hallmark of MMD.^20^. Therefore, protocols for constructing animal models by surgical ligation of arteries are readily available. Choi et al. created a chronic hypoperfusion model of MMD in rats through ligation of bilateral common carotid arteries and found that the impaired function of endothelial colony-forming cells which might play a role in the pathogenesis of MMD^7^. However, the absence of microenvironmental and genetic factors and the unexpected acute injury or even death of animals caused by surgery limit the capability of surgical models to explore mechanisms of MMD.

In addition to in vivo models, the in vitro models also can contribute to exploring the pathogenesis of MMD. In our previous study, we discovered that the upregulation of cytoskeletal proteins in MMD was associated with the intimal hyperplasia of affected blood vessels through a proteomic analysis involving 60 MMD patients and 20 healthy controls, as well as in vitro experiments that utilized the superficial temporal arteries and middle cerebral arteries obtained from MMD patients^17^. This study further elucidated the pathophysiological mechanisms of MMD that threonine kinase/glycogen synthase kinase 3 beta/beta-catenin signaling pathway might play a major role in these abnormal changes in endothelial cells^17^. These findings not only suggested that biological markers in our peripheral blood may be applied to the diagnosis, but also suggested that these markers might be the potential targets for the treatment of MMD. Recently, Organoids have been applied to construct disease research models^11^. Hereditary hemorrhagic telangiectasis (HHT), a genetic vascular disorder, is characterized by vascular dysplasia resulting from enlarged and dilated capillaries lacking pericytes and smooth muscle cells (SMCs) coverage^21^. Koh et al. constructed organoid models of HHT by genome editing and inducing pluripotent stem cell differentiation^21^. They have succeeded in discovering that endothelial cells hypofunction in a 2D culture system of HHT, and reproducing the characteristic vascular changes of HHT in a 3D blood-vessel organoid system^21^. The results of this study suggested that organoid models can play an important role in studying multiple dimensions of disease, including etiology, pathophysiology, diagnosis and treatment.

The α-tubulin encoded by the TUBA gene binds to β-tubulin to form heterodimers, which further polymerize to form microtubules, a key structural component of the cytoskeleton. Microtubules provide scaffolds within cells, help maintain cell shape, and are involved in intracellular material transport, chromosome separation, and cell division. This study suggests that overexpression of TUBA4A may promote the proliferation and migration of vascular smooth muscle cells, which play an important role in the occurrence and development of vascular diseases.

### Limitations of the study

Although organoid models have significant advantages in simulating the physiological and pathological features of blood vessels, they are still not fully representative of the complex pathological environment in vivo. Organoid models may lack certain systemic factors, such as the influence of the immune system, which may affect the interpretation of the results. In addition, a further exploration method can be to transplant vascular organoids into immunodeficient mice, which will allow human and mouse blood vessels to anastomose and reconstruct blood flow in human capillaries to further judge the condition of the model.

### Conclusion

This study successfully established a vascular organoid model derived from patient-induced pluripotent stem cells (iPSCs) and, through single-cell sequencing and proteomics analysis, revealed significant abnormalities in the quantity and function of vascular smooth muscle cells (VSMCs) in Moyamoya disease (MMD). The findings demonstrate that the abnormal overexpression of TUBA4A and TUBB4B may promote phenotypic switching, proliferation, and migration of VSMCs by regulating the GJA1 signaling pathway, ultimately contributing to the intimal thickening observed in MMD, a key pathological feature of the disease.

Our results provide new insights into the pathogenesis of Moyamoya disease and suggest that TUBA4A and TUBB4B could serve as potential therapeutic targets. Future studies should further validate the functional roles of these genes and the underlying pathways and explore their impact in different populations and subtypes to advance the development of novel therapeutic strategies for MMD.

## Supporting information

Supplementary materials provide detailed statistical information about this study.

## Acknowledgement

Thanks to the health care workers in the department for their help with the project. Thanks to all participants for their support and cooperation.

## Authors’ contributions

SH, YL, RW provide ideas and make important contributions in various aspects. JZ, XL, ZY, YR, XY, ZQ, XK, collected clinical sample specimens and information. SH, JZ and XL completed the in vitro part of the experiment. SH, JZ, XL did a lot of single-cell sequencing. ZQ, DD assist organoid culture. SH, ZY, SQ completed the data integration and analysis. All the authors have read and approved the final manuscript.

## Data availability statement

The data presented in the current study are available from the corresponding author upon reasonable request.

