## Supplementary materials provide detailed statistical information about this study. for "Organoid Modeling and Single-Cell Profiling Uncover the Migration Mechanism of Smooth Muscle Cells in Moyamoya Disease"

### Methods

#### Plasmid construction and small hairpin RNA transfection

Plasmids encoding TUBA4A, TUBB4B and GJA1 were purchased from the Boen Company (Guangzhou, China). ShRNAs against UBA4A, TUBB4B and GJA1, and the corresponding controls were obtained from Boen Company (Guangzhou, China). Specific primers were designed targeting the TUBA and TUBB sequences. Based on the multiple cloning sites of the pcDNA3.1 vector, Xho I (CTCGAG) and Xba I (TCTAGA) restriction sites were added to the upstream and downstream of the primers, respectively. The PCR product containing the Xho I and Xba I sites was then purified and recovered. The CDS sequences of human TUBA4A and TUBB4B were amplified by PCR using the following primers: TUBA4A forward, 5′-CCGCTCGAGTGAGACCTGTCACCCCGACT-3′; TUBA4A reverse, 5′-TGCTCTAGATAATGGCACAGCCCCAGCTC -3′; TUBB4B forward, 5′-CCGCTCGAGGTTTGCACCTCGCTGCTCCA -3′; and TUBB4B reverse, 5′-TGCTCTAGAGCTCTTGGGGCGATGTCATC -3′. As for GJA1 plasmids, based on the multiple cloning sites of the pLVX-Puro vector, Xho I (CTCGAG) and BamH I (GGATCC) restriction sites were added to the upstream and downstream of the primers, respectively. The PCR product containing the Xho I and BamH I sites was then purified and recovered. The CDS sequences of human GJA1 were amplified by PCR using the following primers: GJA1 forward, 5′-

CCGCTCGAGTTTCATTAGGGGGAAGGCGT-3′; GJA1 reverse, 5′-CGCGGATCCCTCCAGAACACATGATCTGATGG -3′.

ShRNA plasmids for targeted silencing of TUBA4A (sh-FLNA), TUBB4B (sh-ZYX) and GJA1 (sh-GJA1), and the control non-targeting plasmid (sh-NC) were constructed by inserting the following short hairpin sequences into the PLVX-Puro vector: 5′-CCGGGCTCTCTGTTGACTATGGCAACTCGAGTTGCCATAGTCAACAGAGAGCTTTTTG - 3′ for sh-TUBA4A, 5′- CCGGCGCATCTCTGTGTACTACAATCT CGAGATTGTAGTACACAGAGATGCGTTTTTG -3′ for sh-TUBB4B, 5′-CCGGGCCCAAACTGATGGTGTCAATCTCGAGATTGACACCATCAGTTTGGGCTTTTTG - 3′ for sh-GJA1, and 5′-CCGGCAACAAGATGAAGAGCACCAAC TCGAGTTGGTGCTCTTCATCTTGTTGTTTTTG -3′ for sh-NC. TUBA4A overexpression, TUBB4B overexpression, GJA1 overexpression, sh-TUBA4A, sh-TUBB4B, sh-GJA1 and sh-NC plasmids were transfected into 293 T cells. The supernatants were collected and concentrated to construct the lentiviruses. HBVSMCs cells were transfected with various types of lentiviruses at a multiplicity of infection of 10 and 5 μg/ml puromycin was used for selection for 2 weeks to obtain stably transfected cell lines. PCR and western blotting assays were used to verify the transfection efficiency of the lentiviruses.

**Table S1:** Clinical characteristics and sample information of the patients with MMD.

| No. of Patients | Groups | Sex | Age (years) | Hypertension | Diabetes | Coronary heart disease | Hyperlipidemia | Smoking history | Alcohol taking | Duration of symptoms (months) | Suzuki stage | Usage (Sample) |
|---|---|---|---|---|---|---|---|---|---|---|---|---|
| P1 | IS | F | 34 | NO | NO | NO | NO | NO | NO | 7 | L3/R4 | Organoid |
| P2 | IS | M | 45 | NO | NO | NO | NO | NO | NO | 6 | L5/R1 | Organoid |
| P3 | HEM | F | 41 | NO | NO | NO | NO | NO | NO | 24 | L2/R4 | Organoid |
| P4 | IS | M | 28 | NO | NO | NO | NO | NO | NO | 6 | L3/R4 | Organoid |
| P5 | HEM | M | 36 | NO | NO | NO | NO | NO | NO | 12 | L3/R0 | Organoid |
| P6 | HEM | F | 32 | NO | NO | NO | NO | NO | NO | 16 | L1/R4 | Organoid |
| P7 | HEM | M | 17 | NO | NO | NO | NO | NO | NO | 42 | L4/R5 | DIA (Serum) |
| P8 | HEM | M | 23 | NO | NO | NO | NO | NO | NO | 1.5 | L1/R2 | DIA (Serum) |
| P9 | HEM | M | 27 | NO | NO | NO | NO | NO | NO | 5 | L2/R3 | DIA, Vitro experiment (Serum) |
| P10 | HEM | M | 28 | YES | NO | NO | NO | YES | NO | 4 | L3/R2 | DIA (Serum) |
| P11 | HEM | M | 32 | NO | NO | NO | NO | NO | NO | 6 | L0/R2 | DIA, Vitro experiment (Serum) |
| P12 | HEM | M | 32 | NO | NO | NO | NO | NO | NO | 5 | L3/R3 | DIA (Serum) |
| P13 | HEM | M | 44 | YES | NO | NO | NO | NO | NO | 4 | L3/R1 | DIA (Serum) |
| P14 | HEM | M | 52 | NO | NO | NO | NO | NO | NO | 14 | L3/R0 | DIA, Vitro experiment (Serum) |
| P15 | HEM | M | 40 | NO | NO | NO | NO | NO | NO | 24 | L2/R2 | DIA (Serum) |
| P16 | HEM | M | 40 | NO | NO | NO | NO | YES | YES | 2 | L3/R3 | DIA, Vitro experiment (Serum) |
| P17 | HEM | F | 16 | YES | NO | NO | NO | NO | NO | 12 | L2/R4 | DIA (Serum) |
| P18 | HEM | F | 19 | NO | NO | NO | NO | NO | NO | 4 | L1/R1 | DIA (Serum) |

| | | | | | | | | | | | | |
|---|---|---|---|---|---|---|---|---|---|---|---|---|
| P19 | HEM | F | 26 | NO | NO | NO | NO | NO | NO | 4 | L2/R1 | DIA (Serum) |
| P20 | HEM | F | 34 | YES | NO | NO | NO | NO | NO | 8 | L4/R3 | DIA (Serum) |
| P21 | HEM | F | 35 | NO | NO | NO | NO | NO | NO | 6 | L5/R4 | DIA, Vitro experiment (Serum) |
| P22 | HEM | F | 38 | NO | NO | NO | NO | NO | NO | 11 | L3/R6 | DIA (Serum) |
| P23 | HEM | F | 42 | NO | NO | NO | NO | NO | NO | 5 | L3/R3 | DIA, Vitro experiment (Serum) |
| P24 | HEM | F | 43 | NO | NO | NO | NO | NO | NO | 2 | L2/R3 | DIA, Vitro experiment (Serum) |
| P25 | HEM | F | 43 | NO | NO | NO | NO | NO | NO | 5 | L5/R1 | DIA (Serum) |
| P26 | HEM | F | 46 | NO | NO | NO | NO | NO | NO | 120 | L3/R2 | DIA (Serum) |
| P27 | IS | M | 15 | NO | NO | NO | NO | NO | NO | 144 | L3/R5 | DIA, Vitro experiment (Serum) |
| P28 | IS | M | 20 | NO | NO | NO | NO | NO | NO | 3 | L2/R4 | DIA, Vitro experiment (Serum) |
| P29 | IS | M | 31 | NO | NO | NO | NO | NO | YES | 24 | L0/R2 | DIA (Serum) |
| P30 | IS | M | 34 | NO | NO | NO | NO | YES | NO | 1 | L2/R3 | DIA (Serum) |
| P31 | IS | M | 31 | NO | NO | NO | NO | NO | NO | 12 | L3/R3 | DIA (Serum) |
| P32 | IS | M | 35 | NO | NO | NO | NO | NO | NO | 5 | L4/R2 | DIA (Serum) |
| P33 | IS | M | 38 | NO | NO | NO | NO | YES | NO | 2 | L2/R3 | DIA, Vitro experiment (Serum) |
| P34 | IS | M | 38 | NO | NO | NO | NO | NO | NO | 21 | L5/R2 | DIA (Serum) |
| P35 | IS | M | 38 | NO | NO | NO | NO | NO | NO | 3.5 | L5/R3 | DIA, Vitro experiment (Serum) |
| P36 | IS | F | 42 | NO | NO | NO | NO | NO | NO | 72 | L5/R3 | DIA (Serum) |
| P37 | IS | F | 19 | NO | NO | NO | NO | NO | NO | 1 | L2/R3 | DIA (Serum) |

| | | | | | | | | | | | | |
|---|---|---|---|---|---|---|---|---|---|---|---|---|
| P38 | IS | F | 31 | NO | NO | NO | NO | NO | NO | 3 | L2/R3 | DIA (Serum) |
| P39 | IS | F | 32 | YES | NO | NO | NO | NO | NO | 18 | L2/R2 | DIA, Vitro experiment (Serum) |
| P40 | IS | F | 34 | NO | NO | NO | NO | NO | NO | 4 | L2/R3 | DIA (Serum) |
| P41 | IS | F | 34 | YES | NO | NO | NO | NO | NO | 2 | L3/R3 | DIA (Serum) |
| P42 | IS | F | 35 | NO | NO | NO | NO | NO | NO | 12 | L1/R4 | DIA (Serum) |
| P43 | IS | F | 38 | NO | NO | NO | NO | NO | NO | 48 | L2/R2 | DIA (Serum) |
| P44 | IS | F | 39 | NO | NO | NO | NO | NO | NO | 72 | L4/R3 | DIA, Vitro experiment (Serum) |
| P45 | IS | F | 43 | NO | NO | NO | NO | NO | NO | 3 | L3/R2 | DIA (Serum) |
| P46 | IS | F | 30 | NO | NO | NO | NO | NO | NO | 24 | L1/R3 | DIA (Serum) |
| P47 | HEM | F | 32 | NO | NO | NO | NO | NO | NO | 84 | L5/R3 | ELISA (Serum) |
| P48 | HEM | F | 33 | NO | NO | NO | NO | NO | NO | 3 | L3/R4 | ELISA (Serum) |
| P49 | HEM | F | 46 | NO | NO | NO | NO | NO | NO | 120 | L4/R4 | ELISA (Serum) |
| P50 | HEM | F | 36 | NO | NO | NO | NO | NO | NO | 18 | L5/R4 | ELISA (Serum) |
| P51 | HEM | M | 44 | NO | NO | NO | NO | NO | NO | 36 | L1/R3 | ELISA (Serum) |
| P52 | HEM | M | 34 | NO | NO | NO | NO | NO | NO | 8 | L3/R3 | ELISA (Serum) |
| P53 | HEM | F | 48 | NO | NO | NO | NO | NO | NO | 6 | L5/R3 | ELISA (Serum) |
| P54 | HEM | F | 49 | NO | NO | NO | NO | NO | NO | 6 | L4/R3 | ELISA (Serum) |
| P55 | HEM | F | 50 | NO | NO | NO | NO | NO | NO | 9 | L3/R2 | ELISA (Serum) |
| P56 | HEM | F | 50 | NO | NO | NO | NO | NO | NO | 8 | L3/R3 | ELISA (Serum) |
| P57 | HEM | F | 50 | NO | NO | NO | NO | NO | NO | 14 | L3/R1 | ELISA (Serum) |
| P58 | HEM | F | 54 | NO | NO | NO | NO | NO | NO | 9 | L3/R1 | ELISA (Serum) |
| P59 | HEM | F | 60 | NO | NO | NO | NO | NO | NO | 3 | L3/R3 | ELISA (Serum) |
| P60 | HEM | M | 55 | NO | NO | NO | NO | YES | 5 | 6 | L2/R3 | ELISA (Serum) |
| P61 | HEM | F | 55 | NO | NO | NO | NO | NO | NO | 24 | L5/R0 | ELISA (Serum) |

| | | | | | | | | | | | | |
|---|---|---|---|---|---|---|---|---|---|---|---|---|
| P62 | IS | M | 31 | NO | NO | NO | NO | NO | NO | 1.5 | L1/R1 | ELISA (Serum) |
| P63 | IS | M | 50 | YES | NO | NO | NO | YES | NO | 12 | L3/R3 | ELISA (Serum) |
| P64 | IS | F | 35 | NO | NO | NO | NO | NO | NO | 5 | L0/R2 | ELISA (Serum) |
| P65 | IS | F | 49 | NO | NO | NO | NO | NO | NO | 60 | L3/R5 | ELISA (Serum) |
| P66 | IS | M | 35 | YES | NO | NO | NO | NO | NO | 24 | L1/R1 | ELISA (Serum) |
| P67 | IS | M | 37 | NO | NO | NO | NO | YES | YES | 96 | L3/R2 | ELISA (Serum) |
| P68 | IS | M | 43 | YES | NO | NO | NO | NO | NO | 2 | L1/R3 | ELISA (Serum) |
| P69 | IS | F | 43 | NO | NO | NO | NO | NO | NO | 10 | L3/R3 | ELISA (Serum) |
| P70 | IS | F | 43 | NO | NO | NO | NO | NO | NO | 5 | L3/R2 | ELISA (Serum) |
| P71 | IS | M | 46 | YES | NO | NO | NO | YES | NO | 4 | L0/R3 | ELISA (Serum) |
| P72 | IS | M | 45 | NO | NO | NO | NO | NO | NO | 24 | L3/R3 | ELISA (Serum) |
| P73 | IS | F | 51 | NO | NO | NO | NO | NO | NO | 12 | L3/R3 | ELISA (Serum) |
| P74 | IS | M | 46 | NO | NO | NO | NO | YES | NO | 12 | L3/R3 | ELISA (Serum) |
| P75 | IS | M | 46 | NO | NO | NO | NO | YES | NO | 3 | L1/R2 | ELISA (Serum) |
| P76 | IS | F | 48 | NO | NO | NO | NO | NO | NO | 24 | L1/R2 | ELISA (Serum) |

MMD, moyamoya disease. HEM, hemorrhage; IS, ischemia. M, male; F, female. L, left; R, right. DIA, data-independent acquisition mass spectrometry; ELISA, enzyme-linked immunosorbent assays; Duration of symptoms indicates the duration from the first symptom until the hospitalization.

**Table S2:** Antibodies used in western blot and immunofluorescence staining.

| Antibodies | Dilution (application) | Catalogs |
|---|---|---|
| GAPDH | 1:1000 (WB) | ab8245, Abcam |
| CNN1 | 1:1000 (WB) | 17819, CST |
| SMMHCs | 1:1000 (WB) | ab133567, Abcam |
| S100A4 | 1:1000 (WB) | ab197896, Abcam |
| ZO-2 | 2 μg/mL (WB) | 71-1400, Thermo Fisher Scientific |
| TUBA4A | 1:500 (WB) | DF14454, Affinity |
| TUBB4B | 1:1000(WB) | BF0716, Affinity |
| GJA1 | 1:2000 (WB) | ab11370, Abcam |
| PI3K | 1:500 (WB) | AF6241, Affinity |
| P-PI3K | 1:500 (WB) | AF4372, Affinity |
| AKT | 1:1000 (WB) | 9272, CST |
| P-AKT | 1:2000 (WB) | 4060, CST |
| KLF4 | 1:1000 (WB) | 4038, CST |
| Rabbit Anti-Mouse IgG H&L (HRP) | 1:10000 (WB) | ab6728, Abcam |
| Goat Anti-Rabbit IgG H&L (HRP) | 1:10000 (WB) | ab150083, Abcam |
| αSMA | 1:1000 (IF) | ab215912, Abcam |
| CD31 | 1:500 (IF) | ab202368, Abcam |

**Figure S1: The expression level of smooth muscle cell phenotypic switching markers and pathway proteins in vascular organoids. A,** Western blot showing the expression of smooth muscle cell phenotypic switching markers and pathway proteins in vascular organoids. The bar chart of **B-K** shows the expression level of these proteins.

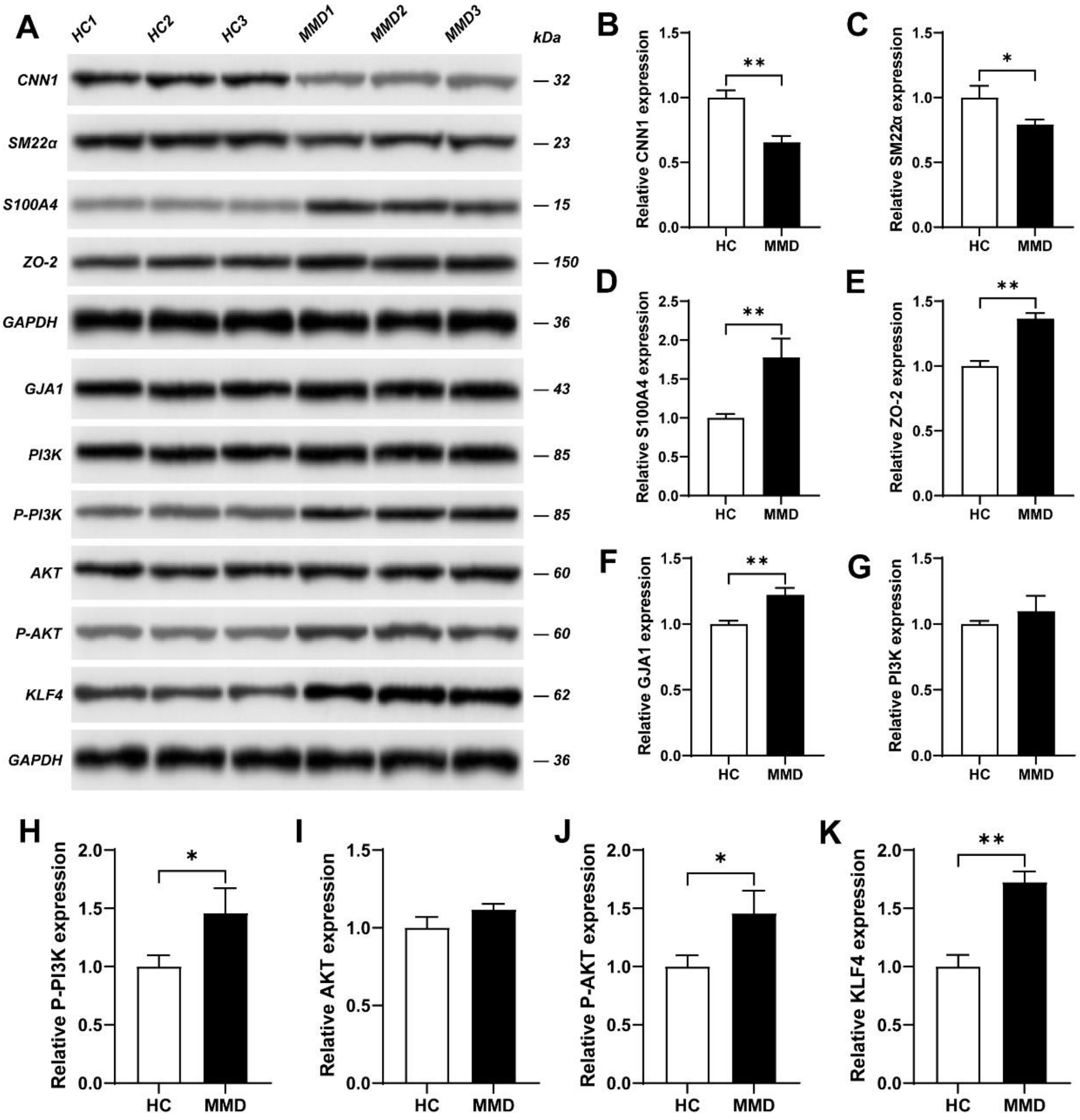

**Figure S2: DIA proteomic detection for the MMD serum. A,** Heatmap of differentially expressed proteins from Data-Independent Acquisition (DIA) proteomic sequencing of serum from MMD patients. **B,** Volcano plot of differentially expressed proteins from Data-Independent Acquisition (DIA) proteomic sequencing of serum from MMD patients. **C,** Chordal graph showing the connections between differentially expressed proteins and various biological processes.

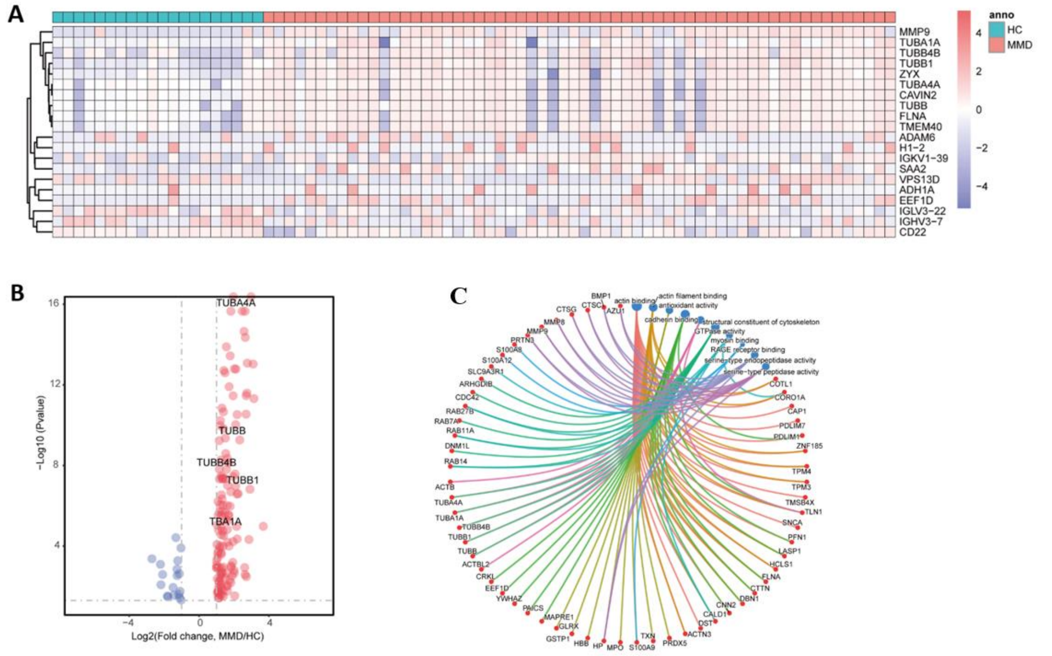

**Figure S3: MMD smooth muscle cells exhibit high abilities of contractile-to-synthetic phenotypic switching, migration and proliferation. A,** Schematic diagram of the establishment for MMD/HC serum-treated HBVSMC cell models. **B,** 5-Ethynyl-2′-deoxyuridine (EDU) staining of the serum-treated HBVSMC. Colors: EDU (red), 4′,6-diamidino-2-phenylindole (DAPI) (blue). Scale = 20 µm. The bar chart of **G** shows the proportion of proliferative HBVSMC. **C,** Scratch assay showing the migration ability of serum-treated HBVSMC. Scale = 200µm. The bar chart of **H** shows the proportion of migration HBVSMC. **D,** Western blot showing the expression level of phenotypic switching markers in serum-treated HBVSMC, including CNN1, SM22α, S100A4, ZO-2. **E,** Western blot showing the expression level of TUBA4A and TUBB4B in serum-treated HBVSMC. **F,** Western blot showing the expression level of GJA1/PI3K/AKT/KLF4 pathway-related proteins in serum-treated HBVSMC.

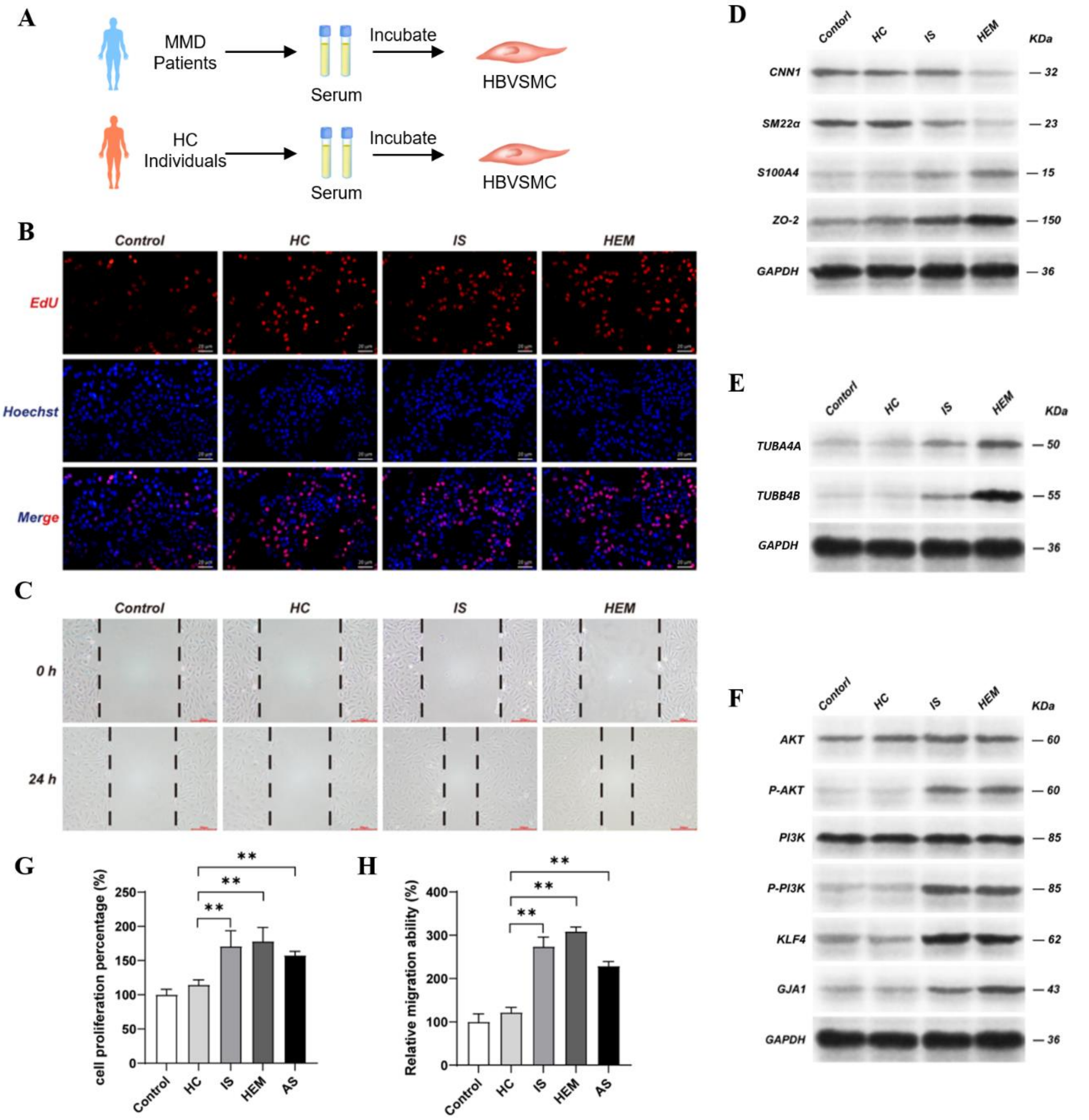


We then used the scratch assay to test the migration ability of HBVSMCs. The results showed that the MMD serum treated HBVSMCs have a higher migration ability compared to the HC serum treated group (**Figure S3C, S3H**). The western blot was used to test the marker of contractile-to-synthetic phenotypic switching. The contractile phenotype biomarkers, CNN1, and SM22α, were downregulated in MMD serum-treated HBVSMCs compared with HC serum-treated HBVSMCs, while the synthetic phenotype biomarkers, S100A4 and ZO-2, were upregulated (**Figure S3D**). Interestingly, we also find the upregulation of TUBA4A and TUBB4B protein in the MMD serum-treated HBVSMC (**Figure S3E**). These results indicates that the MMD serum can promote the abilities of contractile-to-synthetic phenotypic switching, proliferation and migration in HBVSMCs, in which the TUBA4A and TUBB4B protein may play a significant role in this process.

**Figure S4: TUBA4A and TUBB4B promote the abilities of proliferation, migration, and contractile-to-synthetic phenotypic switching of SMC in MMD via GJA1/PI3K/AKT/KLF4 pathway. A,** Western blot analysis showing the efficiency of TUBA4A and TUBB4B knockdown and overexpression. **B,** 5-Ethynyl-2′-deoxyuridine (EDU) staining of the TUBA4A/TUBB4B overexpression HBVSMC. Colors: EDU (red), 4′,6-diamidino-2-phenylindole (DAPI) (blue). Scale = 20 μm. **C,** 5-Ethynyl-2′-deoxyuridine (EDU) staining of the TUBA4A/TUBB4B knockdown HBVSMC. Colors: EDU (red), 4′,6-diamidino-2-phenylindole (DAPI) (blue). Scale = 20 μm. **D,** Western blot analysis showing the efficiency of GJA1 knockdown. **E,** Western blot analysis showing the efficiency of GJA1 overexpression. **F,** Scratch assay showing the migration ability of GJA1 knockdown HBVSMC. Scale = 200μm. **G,** 5-Ethynyl-2′-deoxyuridine (EDU) staining of the GJA1 knockdown HBVSMC. Colors: EDU (red), 4′,6-diamidino-2-phenylindole (DAPI) (blue). Scale = 20 μm. The bar chart of **I** shows the proportion of proliferative HBVSMC. **H,** Western blot showing the expression level of phenotypic switching markers in HBVSMC, including CNN1, SM22α, S100A4, ZO-2.

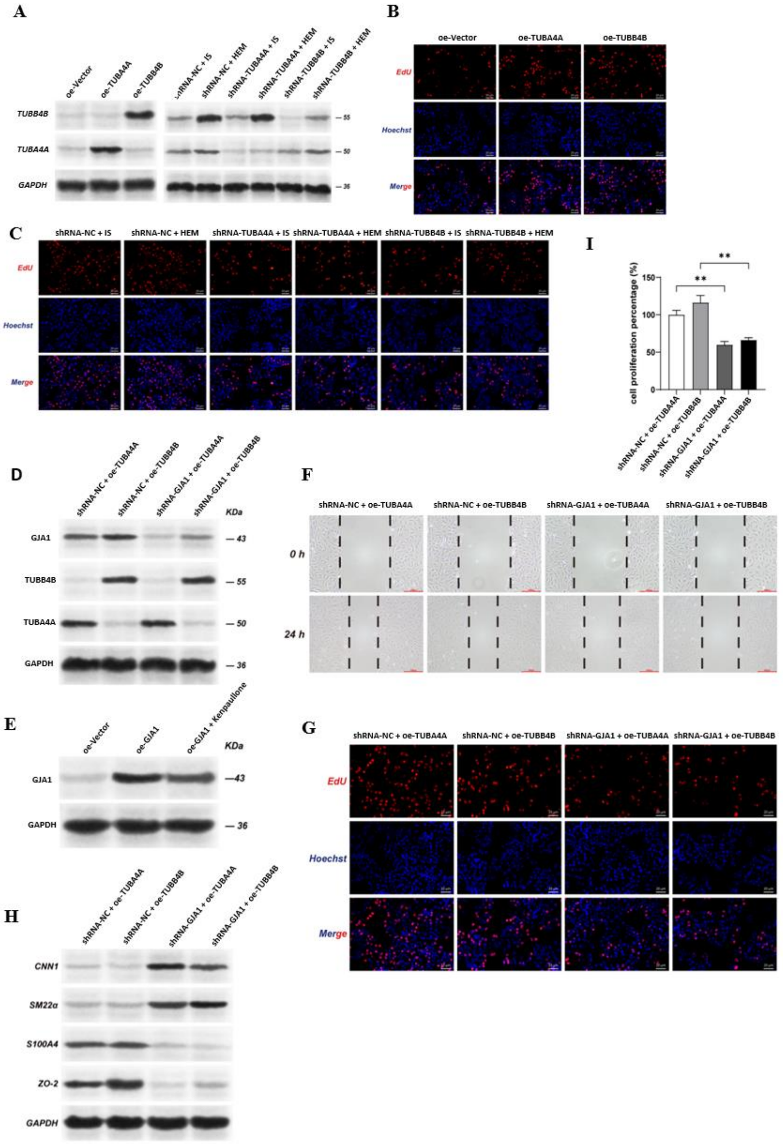

**Figure S5: TUBA4A and TUBB4B promote the abilities of proliferation, migration, and contractile-to-synthetic phenotypic switching of SMC in MMD. A,** Schematic diagram of the establishment of TUBA4A or TUBB4B overexpression and knockdown HBVSMC cell models. **B,** Western blot showing the expression level of phenotypic switching markers in TUBA4A/TUBB4B overexpression and knockdown HBVSMC, including CNN1, SM22α, S100A4, ZO-2. **C,** Scratch assay showing the migration ability of HBVSMC. Scale = 200μm. **D,** Flow cytometric cell cycle analysis of TUBA4A/TUBB4B overexpression and knockdown HBVSMC. The bar chart of **F** and **G** shows the proportion of HBVSMC in G0/G1, S and G2/M cell cycle phases. **E,** Western blot showing the expression level of GJA1/PI3K/AKT/KLF4 pathway-related proteins in TUBA4A/TUBB4B overexpression and knockdown HBVSMC.

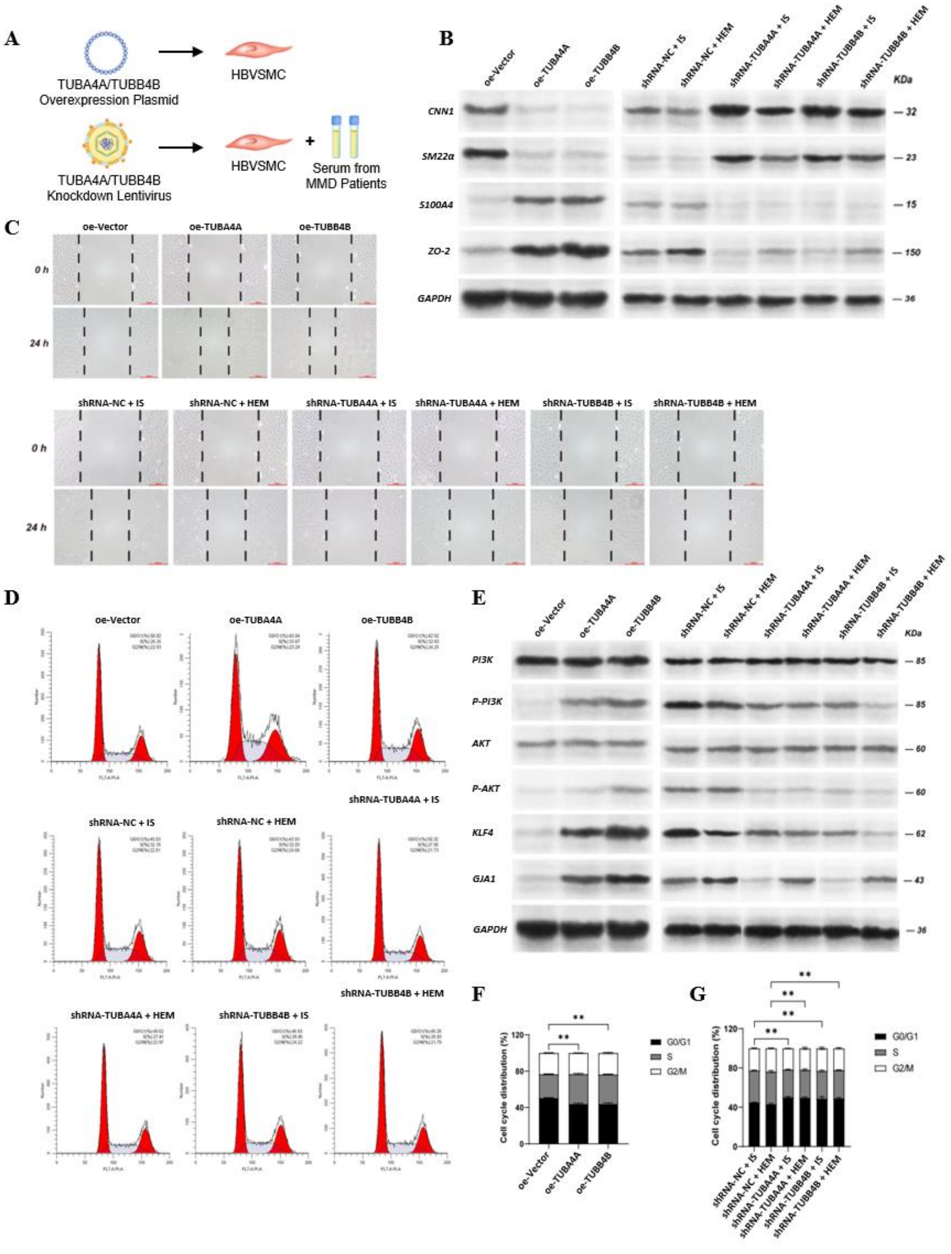


The efficiency of overexpression and knockdown was shown in **Figure S3A**. After the overexpression of the two genes by transfecting oe-TUBA4A and oe-TUBB4B plasmid, the HBVSMCs present a tendency of contractile-to-synthetic phenotypic switching in the TUBA4A and TUBB4B overexpressed group, with the contractile phenotype biomarkers, CNN1, and SM22α, downregulated, and the synthetic phenotype biomarkers, S100A4 and ZO-2, upregulated (**Figure S5B**). The TUBA4A and TUBB4B overexpressed HBVSMCs also have an enhanced ability in cell migration compared with that in the oe-Vector group (**Figure S5C**). The oe-TUBA4A and oe-TUBB4B groups appeared to have more cells in S phase than the oe-Vector group (**Figure S5D, S5F, S5G**). Additionally, the oe-TUBA4A and oe-TUBB4B groups showed a significant increase in VSMC proliferation (**Figure S4G, S4I**). Subsequently, we knocked down these two genes in HBVSMCs, followed by incubation with serum from MMD patients. After 24 hours of serum incubation, the contractile-to-synthetic phenotypic switching after MMD serum treatment was reversed after the knockdown of TUBA4A or TUBB4B (**Figure S5B**). HBVSMCs also exhibited a significantly decreased ability in migration after TUBA4A or TUBB4B knockdown (**Figure S5C**). The TUBA4A or TUBB4B knockdown HBVSMCs showed a reduced proliferation ability (**Figure S5D, S4G, S4I**). Knockdown of the two genes reversed the MMD serum-induced changes of HBVSMCs. These results suggest that TUBA4A and TUBB4B are essential for these changes of HBVSMCs in MMD, including proliferation, migration, and phenotypic switching.

**Figure S6: TUBA4A and TUBB4B induce the changes of VSMC in MMD via GJA1/PI3K/AKT/KLF4 pathway. A,** Schematic diagram of the cell models establishment with GJA1 overexpression, GJA1 knockdown, and KLF4 inhibitor application. **B,** Western blot showing the expression level of GJA1/PI3K/AKT/KLF4 pathway-related proteins in the GJA1 overexpression, GJA1 knockdown, and KLF4 inhibitor applicated HBVSMC. **C,** Western blot showing the expression level of phenotypic switching markers in HBVSMC, including CNN1, SM22α, S100A4, ZO-2. **E,** 5-Ethynyl-2′-deoxyuridine (EDU) staining of the serum-treated HBVSMC. Colors: EDU (red), 4′,6-diamidino-2-phenylindole (DAPI) (blue). Scale = 20 µm. The bar chart of **D** shows the proportion of proliferative HBVSMC. **F,** Scratch assay showing the migration ability of GJA1 overexpression and KLF4 inhibitor applicated HBVSMC. Scale = 200µm.

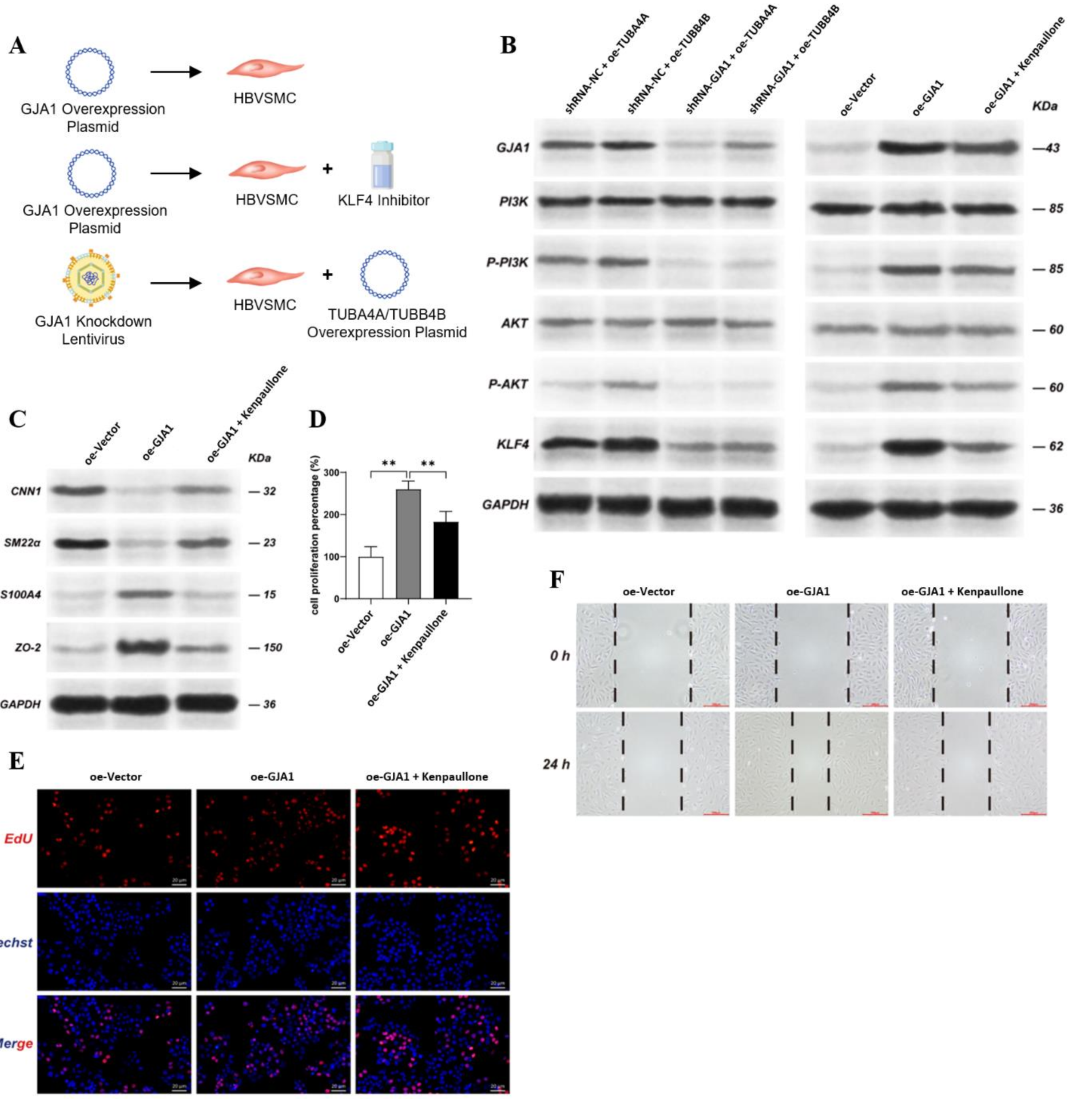


The efficiency of overexpression and knockdown for GJA1 was shown in **Figure S4D and S4E**. The contractile phenotype biomarkers, CNN1, and SM22α, were downregulated in GJA1 overexpressed HBVSMCs, while the synthetic phenotype biomarkers, S100A4 and ZO-2, were upregulated (**Figure S6C**). The GJA1 overexpression group showed an increased proportion of proliferating cells (**Figure S6D, S6E**). Overexpression of GJA1 enhanced the migration ability of HBVSMCs (**Figure S6F**). Conversely, when GJA1 was knocked down in HBVSMCs overexpressing TUBA4A and TUBB4B, we observed these TUBA4A or TUBB4B-induced HBVSMCs changes were reversed (**Figure S4F-I**). These results indicates that the GJA1 plays an important role in these HBVSMCs changes induced by TUBA4A or TUBB4B.

Meanwhile, it was observed that the expressing levels in the PI3K/AKT/KLF4 pathway proteins increased following the overexpression of GJA1, including p-PI3K, p-AKT and KLF4 (**Figure S3F, S5E, S6B**). To investigate whether the PI3K/AKT/KLF4 pathway contributes to the GJA1 overexpressed-caused changes of HBVSMCs, we used Kenpaullone, a KLF4 inhibitor, to downregulate the expression of the downstream KLF4 molecule in this pathway (**Figure S6A**). After the KLF4 inhibition for HBVSMCs, the enhanced proliferation, migration, and contractile-to-synthetic phenotypic switching induced by GJA1 overexpression were all blocked (**Figure S6B-F**). These demonstrate that the GJA1/PI3K/AKT/KLF4 pathway contributes to the TUBA4A and TUBB4B induced changes of HBVSMCs, including contractile-to-synthetic phenotypic switching, and the enhanced abilities of proliferation and migration (**Figure 7C**).
